# Volume and surface methods for microparticle traction force microscopy: a computational and experimental comparison

**DOI:** 10.64898/2026.03.28.714997

**Authors:** Simon Brauburger, Bastian K. Kraus, Tobias Walther, Cornelis Mense, Tobias Abele, Kerstin Göpfrich, Ulrich S. Schwarz

**Affiliations:** Institute for Theoretical Physics, Heidelberg University, 69120 Heidelberg, Germany; BioQuant, Heidelberg University, 69120 Heidelberg, Germany; Cavendish Laboratory, University of Cambridge, CB3 0HE Cambridge, United Kingdom; Center for Molecular Biology of Heidelberg University (ZMBH), Heidelberg University, 69120 Heidelberg, Germany; Cluster of Excellence SynthImmune, Heidelberg University, 69120 Heidelberg, Germany

## Abstract

It is an essential element of mechanobiology to measure the forces of biological cells. In microparticle traction force microscopy, they are inferred from the deformation of elastic microparticles. Two complementary variants have been introduced before: the volume method, which reconstructs surface stresses from the displacements of fiducial markers embedded inside the particles, and the surface method, which infers stresses directly from the deformation of the particle surface. However, a systematic comparison of the two methods has been lacking. Here, we quantitatively compare both approaches using simulated traction fields representing biologically relevant loading scenarios. We find that the surface method consistently reconstructs traction profiles with substantially lower errors than the volume method, which suffers from displacement tracking and stress calculation at the surface. At high noise levels, however, the performance gap becomes smaller. To compare the performance of the two methods in a realistic experimental setting, we developed DNA-based hydrogel microparticles equipped with both fluorescent surface labels and embedded fluorescent nanoparticles, enabling the direct comparison of the two methods within the same system. Compression experiments produced traction profiles consistent with Hertzian contact mechanics and confirmed the trends observed in the simulations. We also show that despite large experimental deformations and strains (both up to 20 percent), linear elasticity theory should still be valid. While our computational workflow establishes a framework to apply both methods, our experimental workflow establishes DNA microparticles as versatile and biocompatible probes for measuring cellular forces.

## 1. INTRODUCTION

The generation and sensing of physical force is an important element in many cellular processes, including cell differentiation, division and migration.^1^ During tissue morphogenesis, for example, cells are guided by the mechanical properties of their surroundings,^2,3^ while immune cells probe mechanical properties to select their targets.^4,5^ To measure the physical forces at work in such biological systems, a large range of different methods has been established in the field of mechanobiology.^6–8^

The standard method to measure forces of single cells is traction force microscopy (TFM).^9,10^ In the classical setup, a single cell is placed on a flat, elastic substrate with embedded fluorescent nanoparticles. The substrate is deformed by the cellular forces, causing measurable displacement of the nanoparticles. By tracking the nanoparticles close to the surface, the displacement field on top of the substrate is obtained, from which the traction field can be reconstructed using elasticity theory. The most efficient method for this purpose is Fourier Transform Traction Cytometry (FTTC), which maximizes the agreement between predicted and measured data (inverse method) and inverts the relation between forces and displacement in Fourier space.^11^ The effect of the inevitable noise in the experimental data can be counteracted by regularization.^12^ Alternatively, one can directly map strain onto stress (direct method), but this requires taking image stacks and derivatives of noisy data.^13–15^. Today, TFM on flat substrates is a very mature method, which has been adapted to different cellular contexts. For both inverse and direct methods, software solutions are available in the public domain.^15–19^

TFM is also routinely applied to multicellular systems. A straightforward example is epithelial monolayers, for which TFM has been extended by monolayer stress microscopy, which predicts stresses in the monolayer from traction forces on soft elastic substrates.^20–22^ This approach can even be applied to intestinal organoids, if they are prepared to adhere to a flat substrate.^23^ To measure cell forces in full three-dimensional contexts, one usually embeds the system of interest in collagen gels.^24,25^ The mechanics of such gels, however, is less well defined, leading to lower resolution.

While all of these methods use spatially extended substrates to measure cell traction, one can also employ small deformable probes embedded into the system of interest. This approach has been pioneered for fluorescently labeled oil droplets.^26^ A huge advantage of this approach is the possibility of these force sensors to be used inside multicellular systems such as embryos, tumor spheroids or organoids.^27,28^ One disadvantage of the oil droplet system is that as a fluid, it can only measure normal and not shear forces.

Over the last years, there has been a growing effort to replace fluid droplets with spherical gels, called microparticles, typically made from polyacrylamide (PAA), the standard material used for spatially extended substrates in TFM.^29–32^ This approach is called *microparticle traction force microscopy* (MP-TFM)^29^ and comes with its own challenges, mainly due to its small system size compared with spatially extended systems. Yet, it is a very appealing approach, because it combines the advantages of traditional TFM and the fluid droplet method: MP-TFM enables the detection of shear forces *in situ*, which means inside living biological systems.

Two main strategies have emerged to reconstruct traction fields from elastic microparticles, as depicted in Fig.1. The first approach, introduced by Mohagheghian et al.,^30^ follows the classical paradigm of traction force microscopy. Fluorescence images of marker nanoparticles embedded throughout the microparticle are recorded both in an undeformed reference state and in a deformed state. From these images, the three-dimensional displacement field 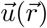 inside the particle volume is obtained using particle tracking or volume correlation algorithms. The traction field is then calculated from the displacement field by evaluating spatial derivatives of 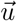 and applying the constitutive relations of linear elasticity. This is known as the direct method in conventional TFM.^15^ In the following, we refer to this variant of MP-TFM as the *volume method*.

**Figure 1.**
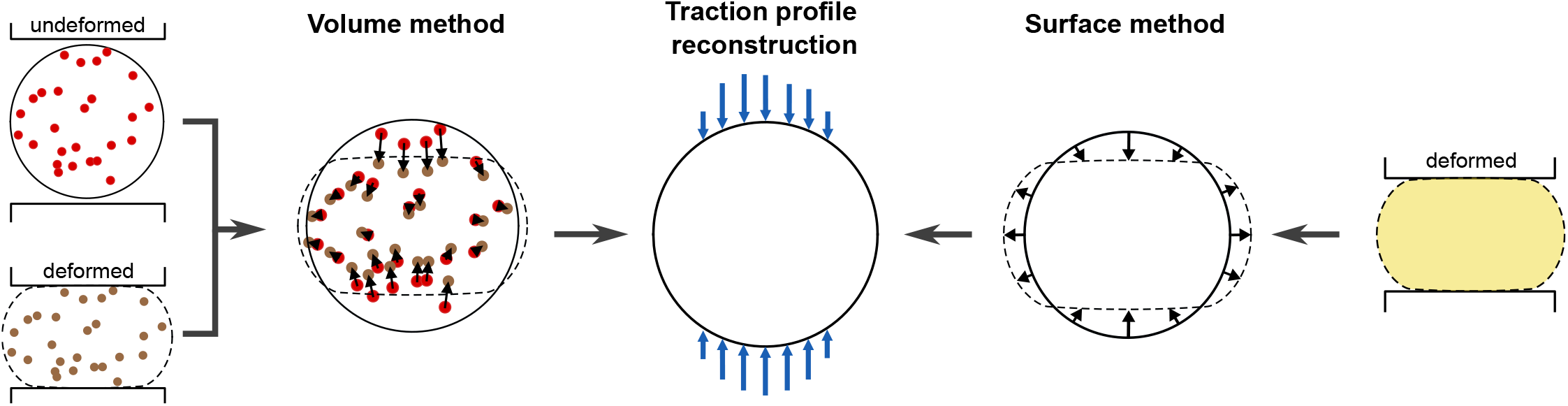
Volume and surface methods in microparticle traction force microscopy (MP-TFM). In the volume method (left), the three-dimensional displacement field is obtained by tracking fluorescent nanoparticles inside the microparticle from the undeformed reference state (red) to the deformed state (brown). Surface tractions are calculated using the direct method, which directly maps strain to stress. In the surface method (right), the geometry of a deformed microparticle (usually homogeneously fluorescent, yellow) is compared to an assumed spherical reference. The traction field is then inferred from the surface displacement using an expansion in vector spherical harmonics. In a simple variant of the surface method, only radial displacements are being considered. Both volume and surface methods ultimately infer a surface traction profile (center), but use different information and assumptions.

The second approach, which we will refer to as the *surface method*, is closer to the fluid droplet method, relies solely on the deformation of the particle surface, and has been pioneered by Vorselen et al.^29,33–35^ From this boundary geometry, surface displacements are inferred relative to an assumed spherical reference configuration. These surface displacements are expanded in vector spherical harmonics, which enable the conversion into a traction field.^31,32,36^ To achieve higher accuracy, the displacement field can then be further refined by iteratively minimizing an appropriate energy functional of the traction and displacement fields, albeit at higher computational cost.^29^ One disadvantage of this method is that the reference configuration is not measured, but assumed. In particular, in the fast variant without refinement, only radial displacements are considered.^31,32^

The two methods differ fundamentally in the type of displacement information they use (volume-versus surface-levels) and in the assumptions they make, especially about the reference configuration. Therefore, it is not evident *a priori* which strategy should provide more accurate or robust traction estimates under realistic experimental conditions. A systematic comparison of both methods has not been conducted before. Here, we present a systematic comparison between the two methods and introduce an experimental system (DNA-based hydrogel microparticles with appropriate fluorescent markers) that allows us to test these two methods also in practice. Although in principle other pipelines might be developed for MP-TFM, e.g., an inverse variant of the volume method, here we restrict our treatment to the two commonly used and already optimized pipelines. This work is structured as follows. First, we computationally compare the accuracy of the two different methods in recovering a known traction profile on the surface of a sphere. For both methods, we simulate suitable experimental data for three different traction profiles mimicking relevant biological scenarios and calculate the errors in recovering the prescribed traction profile. Overall, we find that the surface method achieves lower reconstruction errors than the volume method, making better use of the information about the relevant deformations near the microparticle surface. We attribute this to surface effects of the correlation algorithm used to track the fiducial markers in the volume method, leading to inaccuracies that result in an underestimation of tractions in the area where the force was actually applied. In order to experimentally validate our findings, we then introduce a new experimental system for MP-TFM, namely DNA-based hydrogel microparticles (DNA-HMPs). DNA-HMPs have recently been shown to exhibit elastic solid-like material properties and can be produced in a controllable manner with stiffnesses tunable between 30 Pa and 6 kPa. DNA-HMPs are further easily equipped with guest molecules for chemical stimulation of surrounding cells and stably embedded into organoid and spheroid systems, enabling directed tissue engineering.^37,38^ Employing DNA-HMPs thus offers a biocompatible and tunable approach for 3D force measurements *in vitro*. We test the DNA-HMPs on both the volume and the surface method by compressing them inside a custom-made microwell setup, where the DNA-HMPs are deformed using a weighted glass slide to recreate a Hertzian contact scenario. Both methods retrieve the overall shape of the traction profile. The surface method achieves a closer match to the corresponding analytical solution, in agreement with the simulation results. Finally, we demonstrate through computer simulations with hyperelasticity, that even for the large deformations and strains used in our experiments, linear elasticity is still justified.

## 2. MATERIALS AND METHODS

### 2.1 Volume method

Displacements were estimated using the Fast Iterative Digital Volume Correlation (FIDVC), developed by Franck et al.,^39,40^ and available on GitHub.^41^ FIDVC is based on the Digital Volume Correlation (DVC) algorithm to align two images *I*(*x, y, z*) (reference) and *Î* (*x, y, z*) (deformed), containing gray level intensity values. We first divide the image *I* into subvolumes *V*_*i*_ of integer voxel size lengths *l*_*x*_, *l*_*y*_, *l*_*z*_, also referred to as window size. For each given subvolume *V*_*i*_, we now try to estimate a single displacement vector 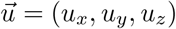 by evaluating a performance metric for different trial displacements 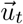 and choosing the one with the best result. As performance metric, we use a cross-correlation function that characterizes the degree of similarity between two pictures by multiplying the intensity values of all voxels in the volume. In our application, these two pictures consist of the reference picture *I* and the deformed picture *Î*, where to the latter 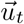 has been applied reversely.

We sum over all the voxels of the subvolume *V*_*i*_:

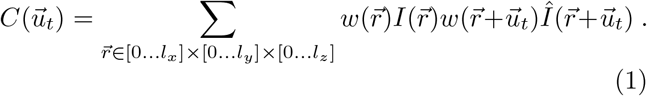

A weight function 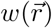 is introduced to compensate for the frequency response of the moving average filter. Furthermore, to increase computational speed and obtain more accurate results, we evaluate Eq.(1) in Fourier space and use a third order polynomial Gaussian peak algorithm.^39^

To decrease computation time, improve resolution, and enable the detection of larger displacements, the procedure described above has been extended into a multi-step iterative algorithm called Fast Iterative DVC (FIDVC).^40^ The main idea is that we do not start with the final resolution, but with a large window size and thus a lower resolution in order to find a coarse grained displacement field. This displacement field is then reversely applied to the images, the window size is halved, and the procedure is repeated until the desired resolution or accuracy is reached.

Once the displacement field has been obtained from the measurement data using the FIDVC, the next step is to calculate the traction from the displacement field. For the volume method, we use the direct method, corresponding to directly mapping displacements to strains and then strains to stresses. The strain tensor describes relative deformations. Assuming small relative deformations, its linear approximation, which significantly simplifies the following calculations, is given via

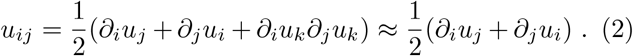

Assuming that our material is an isotropic Hookean solid, we can define the stress tensor that connects the strain and material properties. It is characterized by the Lamé parameters *µ* and *λ*:

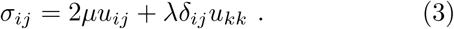

Evaluating the stress tensor across the surface of the body by multiplying it with the normal vector 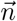 of the surface at that point yields the traction 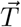, which describes the stresses acting on the surface of the body:

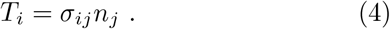

Thus, in order to retrieve the traction, we first need to compute the spatial derivatives of all displacement components to obtain the strain tensor. Each displacement field component as returned from the FIDVC is given as a 3D array with one value for each voxel, so the data is discretized with voxel spacing *d*. This allows us to express the differentiation as a matrix operator, or (finite difference) kernel, performed for each point by the summation of element-wise multiplication of the operator with the surrounding data points. In this work we use the 5-tap kernel:^42^

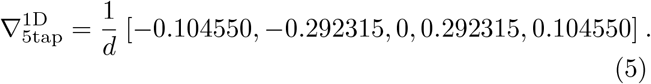

For higher dimensions, the kernel needs to be reshaped for each direction to represent the respective derivative. It is also common to additionally perform averaging over the dimensions orthogonal to the direction of the derivative, resulting in, what is known as, the optimal-5 kernel.^39^

Applying the kernels described above gives us the strain tensor *u*_*ij*_ for each voxel in Cartesian coordinates. In the next step, we calculate the stresses *σ*_*ij*_ using Eq.(2) and Eq.(3) at each voxel. In order to retrieve tractions, we need to evaluate the stresses with the corresponding normal vector at the surface. We use a Gauss-Legendre quadrature (GLQ) mesh as set of points to perform the evaluation (see Fig.S1) because the GLQ mesh is also used extensively in the surface method. Due to the surface effects, we evaluate the traction not on the surface of the sphere, but on a smaller sphere with the evaluation radius *r*_*e*_ *< r*_0_ (more details in Section 3.3.2). We interpolate *σ*_*ij*_ from the surrounding voxels onto the point of the evaluation sphere surface using the RegularGridInterpolator function of numpy. Finally, tractions are obtained with Eq.(4), using the corresponding normal vector of each point on the grid. The traction components, which now represent the magnitude in terms of Cartesian unit vectors, are also transformed to spherical coordinates.

### 2.2. Surface method

Force reconstruction by the surface method is based on minimizing an energy functional which was proposed by Vorselen et al. in 2020,^29^ and uses the decomposition into spherical harmonics introduced by Wang et al. in 2019.^36^ Note that this approach is only possible for finite systems like MPs, so that the surface carries a lot of information. The main idea has some similarity with the inverse method of traditional TFM, because it also minimizes a functional. For MP-TFM, the functional consists of three terms representing various considerations. First, any residual traction on a region which is known to be traction-free is penalized. Second, the elastic energy should be minimal to account for the minimum energy principle. Third, we want to remove aliasing effects by penalizing high-frequency components. As a linear combination, we obtain:

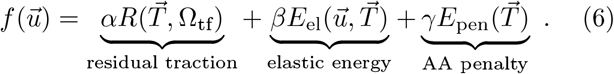

The parameters *α, β*, and *γ* are defined in Table S1; these choices were shown to provide optimal results in the original implementation of the surface method.^29^ To evaluate this functional quickly, it is necessary to use a fast way to convert from 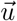 to 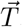, as the terms in the functional have an explicit dependence on 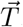 The solution is to choose a basis for the displacement function space where each basis vector 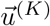 corresponds to a basis vector 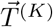. To construct this basis, we start by looking at the space of possible solutions. These need to obey the equilibrium condition

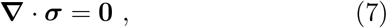

as we assume that the sphere is in mechanical equilibrium. In the next step, we use a harmonic potential ansatz for the potential:^43^

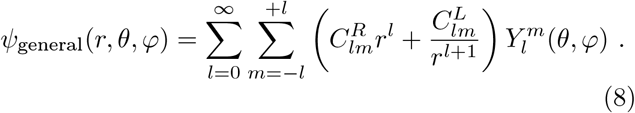

We disregard the terms ~ 1*/r*^*l*+1^, as they diverge at the origin (*r* = 0). The corresponding displacement solutions are then given by the Papkovich-Neuber solutions:^43^

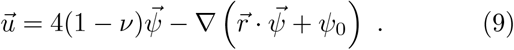

These provide a basis for the relevant displacement solutions, whose angular part is represented as a superposition of spherical harmonics. Evaluating Eq.(2) to Eq.(4), we find a similar decomposition for the traction 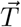. As the decomposition for each basis vector 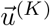 and 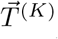 is fixed, the conversion between them reduces to a simple matrix multiplication (for detailed derivation see Note S1^36^). This decomposition enables us to adopt a vector representation of the full displacement field (see Note S2). In this way, we can rewrite the functional and its derivatives in terms of matrices, facilitating a minimization of the functional in a reasonable amount of time. This minimization happens in the subset of displacement functions that match the surface profile (obtained by either experiment or simulation) and is performed using the conjugate-gradient method (detailed explanation in Note S3).

### 2.3. Data simulation

To obtain a reference solution which can be used to simulate experimental data, we use the analytical solutions of axisymmetric stress problems presented by Mietke et al.^44^ This represents a special case of the general solution presented in Section 2.2.2. A given axisymmetric traction profile 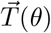 on the surface of a sphere can be decomposed via a set of coefficients *a*_*n*_ and *e*_*n*_, using the orthogonal Legendre polynomials *P*_*n*_:

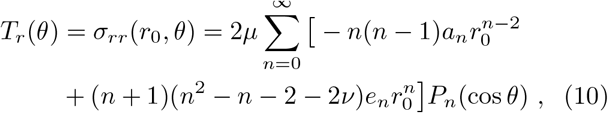

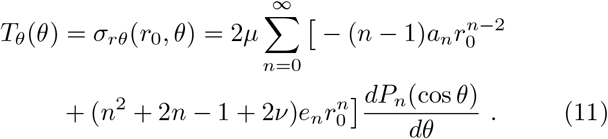

These coefficients directly determine the full solution for the displacement of the sphere resulting from the traction:

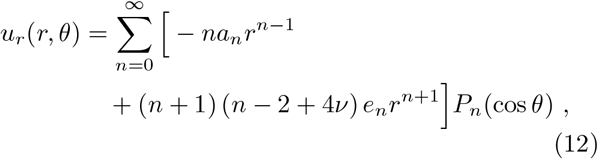

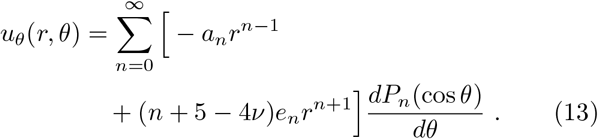

For any boundary condition 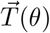, we can now quickly calculate an analytical solution for the full deformation field 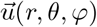 inside the sphere. The described decomposition is of infinite size, so we need to set a cutoff order for the sums. *n*_max_ = 50 was found to be a sufficient compromise between computational speed and accuracy (see Fig.S2).

To simulate image data for the volume method, we create two 3D arrays; *I*(*x, y, z*) for the undeformed and *Î* (*x, y, z*) for the deformed image. Its entries represent intensity values simulating the fluorescence measurements. To obtain these values, the algorithm creates uniformly randomly distributed nanoparticle positions inside a sphere of radius *r*_0_. Each voxel’s intensity value is then generated by evaluating a squared Gaussian point spread function (PSF) for every nearby nanoparticle. For the deformed picture, we apply the deformation field obtained in Eq.(12) and Eq.(13) to each of the nanoparticles and repeat the remaining procedure. In both cases, we add Gaussian displacement noise with amplitude *σ*_disp_ = 0.17 px to the nanoparticle positions before the evaluation of the point spread function, resembling experimentally observed noise (full procedure is explained in more detail in Fig.S3).^45^ If not stated otherwise, we use the simulation parameters as given in Table S1. Resolution as well as microparticle and nanoparticle properties closely resemble the experimental conditions in the original volume method publication.^30^

For the surface method, we generate *N* = 2000 surface points using a Fibonacci spiral lattice, which distributes successive points by the golden angle 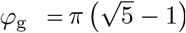. This yields a nearly uniform areal density 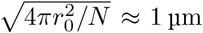, matching the expected resolution of across the entire sphere with a mean surface spacing of the surface reconstructions (a smoothing function with window size 1 µm was used in the original surface method implementation^29^). In contrast to the GLQ grid (used for the actual minimization and efficient traction evaluation by the spherical harmonics approach), the Fibonacci lattice avoids the polar oversampling inherent in latitude-longitude grids, and reconstruction performance between polar and equatorial force application is expected to be unbiased. For each point, we apply the calculated displacement field to obtain the positions of the deformed surface. Noise is implemented as surface roughness, added as a spherical random field in a spectral approach. Random spherical harmonic coefficients *a*_*lm*_ are drawn from a Gaussian distribution with zero mean and a continuous power spectrum

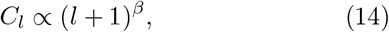

with power-law exponent *β* and minimum degree *l*_min_ = 2 (suppressing monopole and dipole terms to prevent global radius shifts and net displacement offsets). The coefficients are re-normalized so that the root-mean-square (rms) radial perturbation of the resulting field equals a prescribed relative amplitude *σ*_*r*_ (expressed as a fraction of the reference radius *r*_0_). The perturbed surface is obtained by scaling each point’s radial coordinate by (1 + *δ*(*θ, φ*)), where *δ* is the roughness field interpolated at the Fibonacci point locations via bicubic interpolation on the GLQ grid. To characterize the traction-free region, we provide a boolean array. Its entries, each representative for one point, indicate if it is in the traction-free area (True) or not (False). This array is used to calculate the matrix **P** for the functional (see Eq.(33)). Visualizations for an exemplary Fibonacci grid with applied traction field, traction-free regions, and roughness field are given in Fig.S4 and Fig.S5. For the evaluation of the functional, the spherical harmonic cutoff was chosen as *l*_max_ = 20 to maintain feasible runtimes: Changing *l*_max_ from 20 to 30 increased the runtime per period from 5-10 seconds to 90-150 seconds, without a substantial gain in accuracy. We quantify the dependence of the reconstruction quality on *l*_max_ in Fig.S6, confirming *l*_max_ = 20 as a reasonable choice. In this case, there is effectively no need to include the anti-aliasing term in the functional, which only applies for SH components with *l >* 20; for the sake of completeness we still included it in the surface method section. For very localized profiles, it is necessary to increase *l*_max_, depending on the area of force application, see discussion. If not stated otherwise, the simulation parameters as given in Table S1 are used. The same material parameters as for the volume method were chosen to ensure comparability.

### 2.4. Error calculation

To quantify the accuracy of the recovered traction profiles, we assess the normalized mean absolute error (NMAE). In our analysis pipeline, the traction data is evaluated on a GLQ grid with points *x*_*i*_(*θ*_*i*_, *φ*_*i*_) for both surface and volume method. Choosing *θ*_*i*_ ∈ [0, 180°], the NMAE for a traction component is then given by

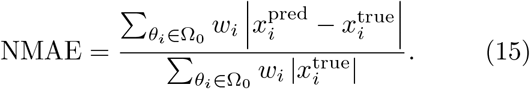

The weights *w*_*i*_ = sin *θ*_*i*_ correspond to the GLQ weights, accounting for the significantly higher point density near the poles due to the smaller spherical surface element d*A* = sin *θ* d*θ* d*φ*.

This *L*_1_-based error metric was chosen for its increased robustness against local noise-induced fluctuations compared to *L*_2_-based metrics. By performing a global normalization with the mean absolute true traction, rather than pointwise normalization, the metric remains well-defined and avoids artificial amplification of errors in regions where the ground-truth traction approaches zero.

### 2.5. Fabrication of DNA-based hydrogel microparticles

DNA was purchased either from Integrated DNA Technologies (unmodified DNA, purification: standard desalting) or Biomers (modified DNA, purification: HPLC). All DNA strands used to form the DNA-HMPs, apart from fluorophore-labeled strands, were diluted in 10 mM Tris (pH = 8) and 1 mM EDTA (Sigma Life Science) to yield 800 µM stock solutions. Fluorophore-labeled strands were diluted in MilliQ water to yield 800 µM stock solutions.

All DNA sequences are listed in Table S2. The DNA stock solutions were stored at −20 °C.

Annealing of the DNA nanostars (A/B) was achieved by mixing the three respective DNA single-strands A-1, A-2, A-3 or B-1, B-2, B-3 at equimolar ratios resulting in 150 µM solutions of the DNA nanostars A and B. In all experiments, 4 mol% of cyanine-3 (Cy3)-labeled strands (B-1-Cy3) were added for visualization by confocal fluorescence microscopy. Phosphate-buffered saline (PBS; Gibco) solution at a final concentration of 1x as well as a final concentration of 10 mM of MgCl_2_ were added for annealing. The nanostars A and B were annealed in a thermal cycler (BioRad) by heating the samples to 85 °C for 3 min and subsequently cooling to 20 °C at an increment rate of 0.1°C/s. To create DNA-HMPs, the nanostars A and B were then mixed at 30 µM final concentration together with the DNA linker strand at 90 µM concentration in a solution of 1x PBS and 10 mM MgCl_2_. Fluorescent nanoparticles (fluorescein-tagged Fluoresbrite yg microspheres (size: 0.3 µm), Cat. No. 24051, Polysciences) were added at 1 vol% to the above solution to create trackable markers (nanoparticles) inside the DNA-HMPs.

The DNA-HMPs were then formed in a droplet-templated manner following the encapsulation of the above solution into water-in-oil droplets with an oil-shell consisting of 2 wt% perfluoropolyether–polyethylene glycol (PFPE–PEG, RAN Biotechnologies) dissolved in HFE-7500 (Iolitex Ionic Liquids Technologies). Encapsulation was achieved by adding the aqueous solution on top of the oil phase in a volumetric ratio of 1:3 in a microtube (Eppendorf, typically 50 µL aqueous to 150 µL oil phase) and vortexing.^37,38^ The droplets were then stored at 21 °C room temperature for 72 h to allow the DNA-HMPs inside the droplets to fully assemble. After that, the DNA-HMPs were released from the water-in-oil emulsion by first adding the buffer solution of 1x PBS and 10 mM of MgCl_2_ at 1.5x of the initial volume on top of the droplet emulsion, and subsequently breaking the emulsion. To break the droplet emulsion, 1H,1H,2H,2H-Perfluoro-1-octanol (Merck) was added on top of the buffer and droplet emulsion, and the mixture incubated for 30 min. The released DNA-HMPs in solution were then transferred to a new microtube for further use.

To equip the DNA-HMPs with a lipid coating, small unilamellar vesicles (SUVs) were formed. They were produced by mixing lipids (99% 1,2-dioleoyl-sn-glycero-3-phosphocholine (DOPC, Avanti Polar Lipids, Inc.) and 1% Atto-633-labeled 1,2-dioleoyl-sn-glycero-3-phosphoethanolamine (Atto-633-DOPE, Atto-Tec GmbH)) dissolved in chloroform (Honeywell) in a glass vial. The solvent was then removed under nitrogen gas and the glass vial placed into a desiccator for 20 min. The lipids were resuspended at a concentration of 10 mM in 1x PBS (Gibco) by shaking the glass vial at 900 rpm at room temperature for 10 min. From this solution, SUVs were extruded through a polycarbonate filter (100 nm pore size, Whatman) with 21 passages. The extruded SUVs were stored at 4 °C. All lipids were stored at −20 °C in chloroform.

Separately, cholesterol-tagged A-nanostars (A-chol) were prepared by annealing the single strands A-1-chol, A-2 and A-3 at equimolar ratios resulting in a 150 µM solution of A-chol. Prior to mixing, A-1-chol strands were incubated at 60 °C for 1 min to break up any aggregates of the cholesterol. Briefly, 50 µL of a DNA-HMP suspension was centrifuged using a C1008-GE myFUGE mini centrifuge (Benchmark Scientific) for 1 min, removing 30 µL of supernatant after centrifugation. The DNA-HMP pellet was then mixed with a final concentration of 5 µM A-chol and incubated for one hour to ensure uptake of A-chol into the DNA-HMPs. The DNA-HMPs were then washed three times with a 1x PBS and 10 mM MgCl_2_ solution before adding SUVs at a final concentration of 1 mM lipids on top of the DNA-HMPs, incubating at 4 °C overnight. Following three more washing steps after the overnight incubation using 1x PBS and 10 mM MgCl_2_ buffer, the lipid-coated DNA-HMPs were ready to use.

### 2.6. Fabrication of the microwell trapping device

Microwells for DNA microparticle trapping were fabricated using a custom-engineered setup, consisting of a Polygon digital micromirror device (DMD) pattern illuminator (MIGHTEX Polygon 1000) coupled to an inverted fluorescence microscope (Axio Observer 7, Carl Zeiss AG). The system was controlled by custom-written scripts based on Python 3.12.4 and the ZEISS ZEN Macro Environment. Printing was performed with a 5× objective (Plan-Apochromat 5×/0.16 M27, Carl Zeiss AG), defining the field of view for each exposure. Binary illumination masks consisting of square microwell frames and a three-letter alphabetical barcode were generated programmatically. To pattern the full sample area, adjacent 5x tiles were sequentially exposed by looping over the entire probe area, with the barcode letters automatically incremented for each microwell and tile. Illumination was provided by a built-in 385 nm LED, and each tile was exposed for 800 ms. Samples mounted on the microscope consisted of a 3D-printed PLA-based four-well chamber slide glued to a silanized 24 mm x 60 mm x 170 µm glass slide using two-component adhesive (ecosil, picodent). The glass slides were silanized to allow for covalent binding of the printed microwells. For this, glass slides were cleaned by sonication in ethanol for 15 min, plasma-activated for 7 min at 200 W using ambient air (PVA TePla 100, PVA TePla AG), and subsequently immersed in a solution of 2% v/v trimethoxysilylpropyl methacrylate (TMSPMA) in toluene (300 mL) overnight, prior to washing in ethanol and MQ water. The wells were filled with a photoresist comprising 50 v% PEGDA 575 (Sigma Aldrich) in MQ water, 5 g/l lithium phenyl-2,4,6-trimethylbenzoylphosphinate (LAP, Sigma Aldrich), 5 g/l ascorbic acid (Sigma Aldrich), and 10 g/l tartrazine (Sigma Aldrich).

### 2.7. DNA-HMP deformation and imaging

The printed microwells were coated for 5 min with poly(vinyl-alcohol) (PVA, 50 mg*/*mL, Sigma Aldrich). The wells were subsequently washed 3x by rinsing with a buffer solution of 1x PBS and 10 mM MgCl_2_. To soak the wells in the same buffer solution, 200 µL of the buffer remained in the wells after washing. Prior to imaging, 10 µL of the DNA-HMP suspension was evenly added onto the buffer-soaked wells and allowed to settle for 20 min. Next, a likewise PVA-coated glass coverslide (diameter: 10 mm, Carl Roth) was added onto the wells. The DNA-HMPs in the wells were imaged on an LSM 900 Zeiss confocal fluorescence microscope (Carl Zeiss AG). Using a Plan-Apochromat 20x/0.8 Air M27 objective, a tile-scan of the entire area containing the microwells was taken and later used as a reference for DNA-HMP deformation. Suitable DNA-HMPs were then selected and the corresponding positions were saved. Using a 63x/1.2 W korr objective with oil immersion, reference z-stacks of the DNA-HMPs in the microwells were acquired using resolution-optimized Airyscan mode and 3x digital zoom at 512 px xy-resolution with 200 nm z-steps. The factor to convert voxels (pixels) to microns is approximately 0.099 for the x- and y-directions and 0.2 for the z-direction, resulting in an anisotropic resolution that is worse in z by a factor of about 2. To deform the DNA-HMPs, a 3D-printed circular weight (polylactic acid, white, diameter = 9 mm, 0.5 g) was placed on top of the glass coverslide, pushing it down and onto the DNA-HMPs. Next, z-stacks of the deformed DNA-HMPs were collected in the same manner as the reference stacks. Lifting off the adapter, the DNA-HMPs were allowed to relax from their deformed state to an undeformed one over several minutes and imaged again using the same settings, allowing us to observe the DNA-HMPs during relaxation. Mechanical characterizations of the DNA-HMPs were also performed to measure relevant elastic parameters (see Fig.S7).

### 2.8. Preprocessing of DNA-HMP images

To obtain suitable image pairs from experimental data for the correlation algorithm in the volume method workflow, the undeformed and deformed images of the nanoparticle configurations inside the DNA-HMPs were aligned with each other, including centering, cropping, and thresholding steps using custom Python scripts. To recover a set of points describing the microparticle surface for the initial radial displacement field guess for the surface method, the fluorescent information of the Cy3-tagged DNA network was analyzed, including blurring and thresholding in ImageJ,^46^ and surface reconstruction steps in GeoV.^47^ The detailed procedures are explained in Fig.S8 to Fig.S10.

## 3. RESULTS AND DISCUSSION

### 3.1. Performance comparison using simulated traction fields

To systematically compare the surface and volume methods, both were applied to simulated experimental data of three axisymmetric traction profiles, shown in Fig.2. The first profile, known as Hertzian contact, mimics the compression of a sphere between two walls, replicating interactions with large bodies, such as other cells or rigid surfaces (Fig.2(a)). The magnitude of indentation in the Hertzian contact profile is characterized by the contact radius *a*. The overall shape of the relevant traction component *T*_*z*_ is recovered successfully by both methods (Fig.2(b)), even though the surface method matches the original profile more closely. As visible in both the deviation maps (Fig.2(c)) and the super-imposed cross-section (Fig.2(d)), the reconstruction of the volume method appears attenuated (~ 30%) and spatially broadened, underestimating the peak magnitude and spreading significantly into the traction-free region. In contrast, the surface method exhibits a slight overshoot and ringing at the edge of the traction profile, likely due to the Hertzian contact’s singular slope at the contact boundary, which is difficult to resolve with a finite-order spherical harmonic representation. The other two recovered traction components *T*_*x*_ and *T*_*y*_ are shown in Fig.S11.

**Figure 2.**
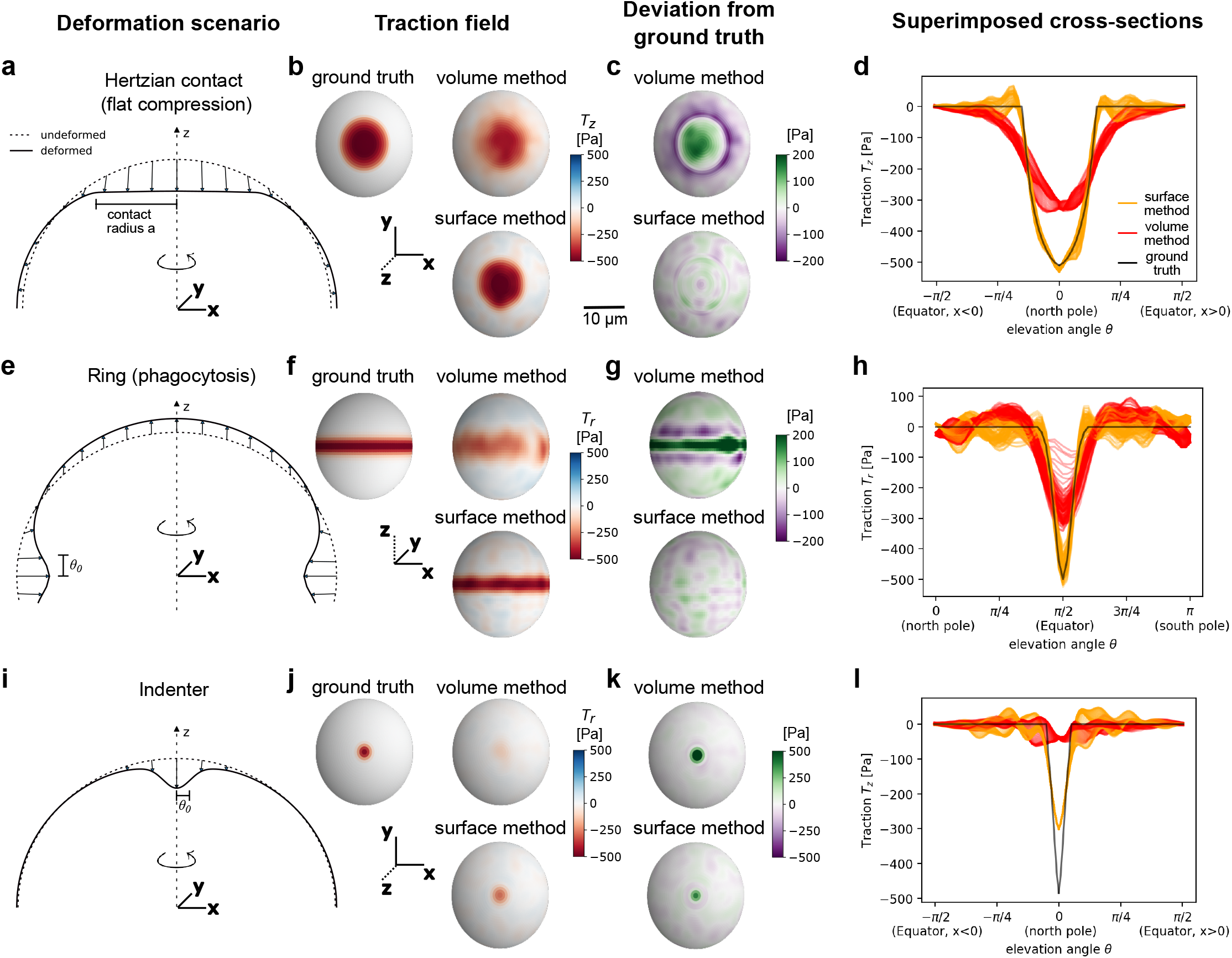
Traction reconstruction for three axisymmetric loading scenarios. (a-d) Hertzian contact (flat compression). (a) Top half of the deformation geometry, which is symmetric around the xy-plane for this and the following two scenarios. (b) Ground truth traction field of the principal traction component *T*_*z*_, followed by reconstructions obtained with the volume and surface methods. The surface method closely reproduces both the magnitude and spatial distribution of the reference traction. The volume method captures the overall structure. (c) Deviation and superimposed cross-sections (d) of the relevant traction component, showing that the volume method exhibits underestimation in the loaded region and compensating deviations in adjacent traction-free areas. (e-h) Ring (phagocytosis-like equatorial indentation). Again, both methods recover the overall profile shape of *T*_*r*_, with the surface method matching the ground truth more closely. (i-l) Indenter. For this strongly localized loading profile, the surface method recovers the spatial profile of *T*_*r*_ but underestimates peak magnitude. The volume method shows pronounced attenuation and spatial spreading of the applied traction. Full traction profiles are provided in Fig.S11 to Fig.S13. Simulation parameters: *E*_0_ = 1500 Pa, *ν* = 0.4, *a* = 0.5 (Hertzian contact), 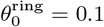, 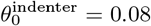, *T*_0_ = 500 Pa (ring and indenter).

The second profile is a Gaussian ring-like indentation, reproducing the compression of a sphere observed during phagocytic engulfment (Fig.2(e)).^29^ The shape of the Gaussian is characterized by *θ*_0_ (width) and *T*_0_ (magnitude), see also Note S4. Again, the recovered *T*_*r*_ component of the volume method appears attenuated (~ 30%) and spatially broadened, albeit less strongly than for the Hertzian contact (Fig.2(f–h)). Interestingly, at the same time the volume method significantly overestimates the magnitude of the *T*_*θ*_ component (~ 5-fold), and also shows spatial broadening for the component (Fig.S12). A potential explanation is that the underlying displacement field is already blurred in the volumetric reconstruction. Because the direct volume method obtains stresses by spatial differentiation of this field, the broadened displacement gradients can produce artificial angular shear, which manifests as a spurious *T*_*θ*_ component. For the surface method, *T*_*θ*_ is also overestimated, but to a lesser extent (~3-fold) and without spatial broadening, likely because the surface-based regularization enforces smoother and more physically consistent traction fields.

The third profile reflects a Gaussian indentation, localized at the poles and designed to assess the reconstruction of point-like traction profiles as observed with focal adhesions or viral/particle entry (Fig.2(i)). Again, the shape of the Gaussian is characterized by *θ*_0_ (width) and *T*_0_ (magnitude). For the width *θ*_0_ = 0.08 examined here, the shape of the volume method reconstruction is barely visible, but resembles the expected shape (Fig.2(i)). The magnitude, however, is strongly attenuated (~ 90%, Fig.2(l)). For the surface method we also observe a significant attenuation (~ 40%), which was not present for the other profiles, and an overestimation of the *T*_*θ*_-component (Fig.S12). The significant attenuation for the surface method can be attributed to the finite spatial resolution imposed by the maximum spherical harmonic order *l*_max_. When the width of the traction profile approaches the corresponding spatial sampling limit, the reconstruction amplitude decreases. This effect is explicitly shown in Fig.S14, where the performance of the surface method as a function of *θ*_0_ significantly drops below a certain width of the indenter.

The closed-form equations of all three traction profiles are shown in Note S4. The simulation and analysis parameters, described in Table S1, are based on experimental studies.^29,30^ The stiffness of the microparticles is characterized by Young’s modulus *E*_0_ and Poisson’s ratio *ν*. The set noise levels led to comparable variations in the vanishing traction components (see *T*_*x*_*/T*_*y*_ in Fig.S11 and *T*_*φ*_ in Fig.S12), enabling a fair comparison.

We then numerically evaluated the errors across the sphere for a range of different noise amplitudes and indentation strengths using the normalized mean absolute error (NMAE) (Fig.3, numeric error values see Fig.S15). Comparing both methods, we find that the errors for the surface method are consistently lower for all examined scenarios. As expected, the performance generally decreases with noise for both methods. Even though the absolute error values of the volume method are higher, the sensitivity of those errors to noise appears to be much lower. This indicates a robustness of the volume method to displacement noise in the microparticles.

**Figure 3.**
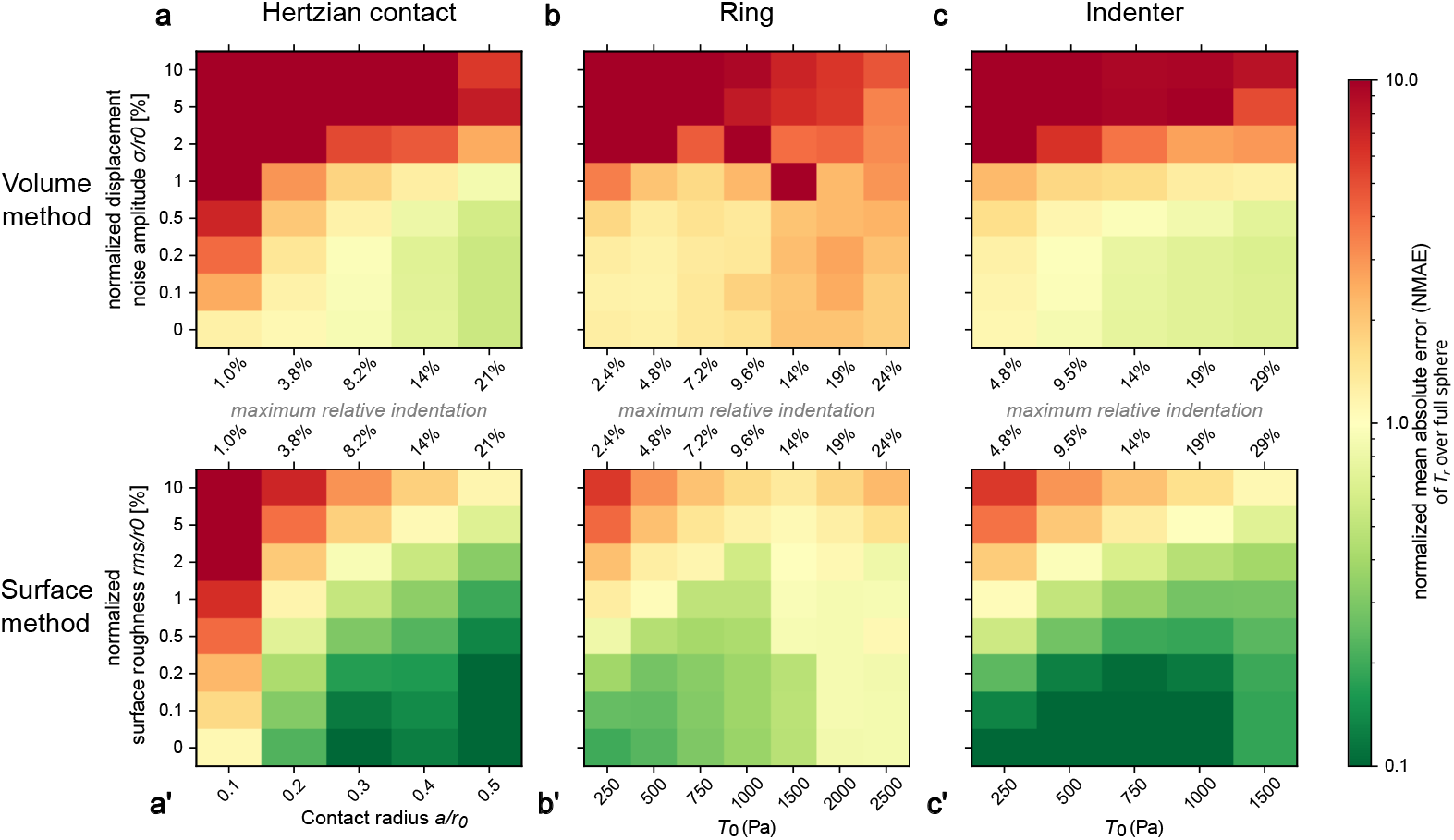
Reconstruction error scaling with displacement noise and loading strength. Normalized mean absolute error (NMAE, evaluated across the full sphere) of the radial traction field component *T*_*r*_ reconstructed by volume method (a) and surface method (b) for the Hertzian contact, ring, and indenter scenarios, evaluated over a range of noise levels and loading parameters. Given the chosen simulation parameters, 1% displacement noise corresponds to one pixel. An extended version of this figure is shown in Fig.S15. Traction profile parameters: *E*_0_ = 1500 Pa, *ν* = 0.4, 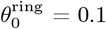, 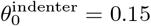, simulation parameters see Table S1.

The volume correlation algorithm appears to recover the field reliably, up to a certain noise level (nanoparticle displacement noise ~ 1 pixel) where the errors diverge. The higher noise sensitivity of the surface method is expected as the surface roughness noise directly propagates into the measured surface displacements and therefore the corresponding traction. For the Hertzian contact profile, the errors of both methods decrease with larger indentation, minimum NMAE error values are 55% (volume) and 8% (surface), respectively, for a maximum relative indentation of 21%. For the ring profile, both methods show a significant decrease in performance with an increase in magnitude of the deformation, increasing from 20% to 87% (surface) and from 120% to 190% (volume) from the lowest to highest indentation considering no noise (maximum relative indentation ranging from 2.4% to 24%). For the indenter, lowest absolute NMAE values across all methods and scenarios of 3% were observed at 10-15% maximum relative indentations with the surface method considering no noise. Here, for the volume method, the lowest error values of 66% were observed at higher magnitudes around 30% indentation, again displaying a remarkable robustness to noise increase. The described trends did not change when the error evaluation was constrained to a localized region around the maximum force application (Fig.S16).

We also evaluated the aforementioned variant of the surface method where a lower number of spherical harmonics is used (*l*_max_ = 5), and only radial displacements are assumed.^31,32^ This corresponds to the initial guess before the optimization of 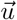 via the functional. This variation of the surface method shows higher errors than both volume and surface method in most cases apart from very high noise and indentation levels, as shown in Fig.S15. When the error evaluation was constrained to a region around the maximum force application (see Fig.S16), this “truncated” surface method outperformed the volume method, but both were again outperformed by the surface method. Following the discussion above and in Fig.S14, this can partially be explained by the lower number of spherical harmonic coefficients, which may not be able to sufficiently resolve the localized nature of the ring and the indenter (effective width ~ 0.2/0.3 rad, minimum effective wavelength of Legendre polynomial ~ 0.6). However, even for the more delocalized Hertzian contact, the errors of the truncated surface method are much higher (e.g., 90% versus 9% of surface method at *a* = 0.3). This indicates that while computationally expensive, the optimization of the displacement solution via the energy functional appears to be the crucial step that gives the surface method its competitive edge. One may argue that this is caused by the significant additional information provided in the traction-free region. However, previous simulations using a scenario comparable to a Hertzian contact with *a* = 0.3 found that leaving out the traction-free region made almost no difference in performance in the magnitude range where the surface method significantly outperforms the truncated one.^29^ A possible explanation for why removing the information about the traction-free regions does not substantially affect the recovery is that the elastic energy and the traction-free penalty term have a similar potential landscape in the functional and minimize 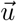 into similar directions. The elastic energy term appears to penalize non-physical displacement profiles. Accordingly, the assumption of the surface displacements being only radial 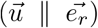 appears to not describe the physical reality of most systems, which is also apparent from considering the deformation sketches in Fig.2(a, e, i). Despite these shortcomings, there are regions of similar performance between surface method and truncated surface method, particularly at higher noise levels, where the truncated approach with significantly lower computational runtimes can be deemed sufficient.

### 3.2. Surface effects of volume method

To understand the substantial performance deficits of the volume method, we examined both its experimental and analysis hyperparameters. First, we systematically varied nanoparticle size and nanoparticle seeding density to assess whether this reduced reconstruction errors, with full results for each configuration given in Fig.S17. While we found that the error does depend on both parameters, and increased significantly especially for small nanoparticle diameters (50 nm) under the resolution limit, the parameters used throughout the simulations (200 nm diameter and 0.002 nanoparticles/voxel) are close to the optimal parameter choices for all scenarios and even the lowest error values obtained are still significantly above those of the surface method, showing that the experimental hyperparameters of the volume method are not responsible for the higher errors.

Next, we examined the behavior of the volume correlation algorithm FIDVC^40^ close to the surface and its connection to the evaluation radius *r*_*e*_, an important analysis parameter. *r*_*e*_ defines the radius of the sphere on which the derivatives of the displacement field are evaluated to obtain the traction profile. In the original volume method publication,^30^ *r*_*e*_ = 0.8 *r*_0_ due to problems with the recovered displacement field directly at the surface. The problematic nature of these “surface effects” can be demonstrated when applying the volume method to the simplest deformation possible: the isotropic compression of a sphere under a given pressure, i.e. only a change in radius. Fig.4(a) shows the values for the radial displacement *u*_*r*_ recovered with the FIDVC for the isotropic compression, which are very accurate up to *r/r*_0_ = 0.8 and then show slight deviations closer to the sphere surface (*r* = *r*_0_), with the absolute values still being reasonably close. However, to calculate the traction, we have to compute the strain *u*_*rr*_ and therefore the derivative ∂_*r*_*u*_*r*_. As a result, what at first sight appears to be a minor absolute deviation, leads to very strong relative deviations near the sphere surface, as shown in Fig.4(b). We also observe stronger deviations in the z-than in the x- and y-directions, which we attribute to the lower resolution we set in that direction to mimic the lower resolution in that direction typical for z-stacks.

**Figure 4.**
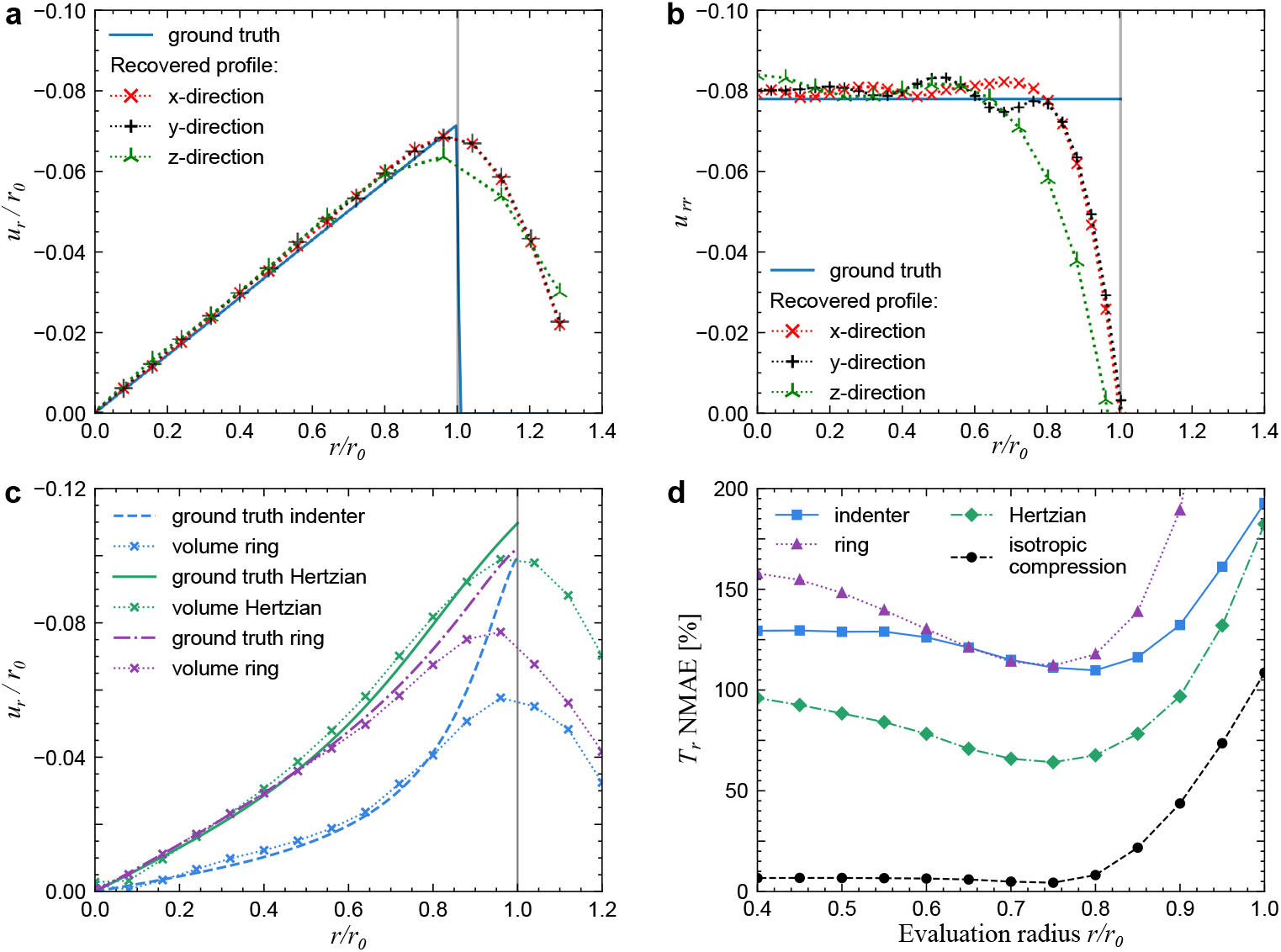
Surface effects of the volume method appear to be the main reason for the higher error rates. (a) Radial displacement *u*_*r*_ for isotropic compression profile. (b) Strain component *u*_*rr*_ = ∂_*r*_*u*_*r*_ (derivative of subfigure (a)) for isotropic compression profile, note that *T*_*r*_ ~ *u*_*rr*_. (c) Radial displacement *u*_*r*_ for the previously examined scenarios, showing even larger deviations near the surface. (d) Normalized mean absolute errors for all scenarios, showing larger errors for localized profiles (ring, indenter).

For the isotropic profile, *u*_*rr*_ and therefore also the traction is constant throughout the sphere due to the constant slope observed in Fig.4(a). Therefore, decreasing *r*_*e*_ still leads to accurate results for this scenario. However, for more complex and especially more localized profiles, the slope changes significantly towards the sphere surface, shown in Fig.4(c). The deviation of the FIDVC results from the ground truth is most prominent for the very localized indenter profile with the steep slope increase near the surface. Therefore, evaluating the components anywhere inside the sphere leads to significant underestimation in the traction profile for the volume method, as evident from Fig.4(d). Regardless of evaluation radius, the obtained errors for the three examined scenarios are all more than a magnitude higher (60-200%) than that obtained for the isotropic compression (*<* 10%). For this reason, the volume method is very suitable for measuring isotropic compression. In its standard configuration without a reference image, the surface method is unable to measure compression, as for the initial guess of the radial displacement solution it assumes an incompressible sphere. In theory, however, pressure could be measured by considering a reference picture of the microparticle and measuring the difference in total volume.

These findings point towards a central problem of the volume method: We can either move the evaluation radius inside to get a more accurate measurement of the displacement field, but lose the important information near the surface, or we move it further to the surface, but then lose accuracy because the volume correlation algorithm is not able to properly track displacements anymore due to the lack of nanoparticles close to and outside of the surface. The decomposition described in Eq.(10) to Eq.(13) explains the underlying reason. When decomposing the traction on the surface via the Legendre polynomials, profiles that are more localized on the surface result in larger coefficient terms *a*_*n*_ and *e*_*n*_ for higher orders of n, corresponding to the necessity of higher frequency components to resolve the profile. These higher order terms then also result in higher powers of *r* for the displacement, considering the scaling of components ~ *a*_*n*_*r*^*n*−1^ and ~ *e*_*n*_*r*^*n*+1^. The higher powers *r*^*n*^ then lead to larger values near the surface of the sphere.

### 3.3. Experimental evaluation using DNA hydrogel microparticles

Finally, we experimentally implemented the volume and surface methods to compare their performance in practice. For this purpose, we employed DNA hydrogel microparticles (DNA-HMPs), which enable both approaches within a single experimental system. As shown in Fig.5(a), the cross-linked DNA forms a fully fluorescent sphere allowing us to recover the sphere surface needed for the surface method, while embedded nanoparticles facilitate the tracking necessary for the volume method. We additionally decorated the DNA-HMP surface with small unilamellar vesicles (SUVs) in an effort to more directly label the DNA-HMP surface. Reconstruction of the DNA-HMP surface from the SUV-coating images, however, proved unsuccessful for our setup (Fig.S10). Nonetheless, SUV-addition remains attractive for future experimental settings in vitro as the SUVs can be used to functionalize the DNA-HMP surface with ligands or receptors, forming more natural interfaces with cells. We also tuned DNA-HMP stiffness to *E*_0_≈1.1 kPa (Fig.S7), which is comparable to values utilized in previous studies (~ 1.4 kPa^30^).

**Figure 5.**
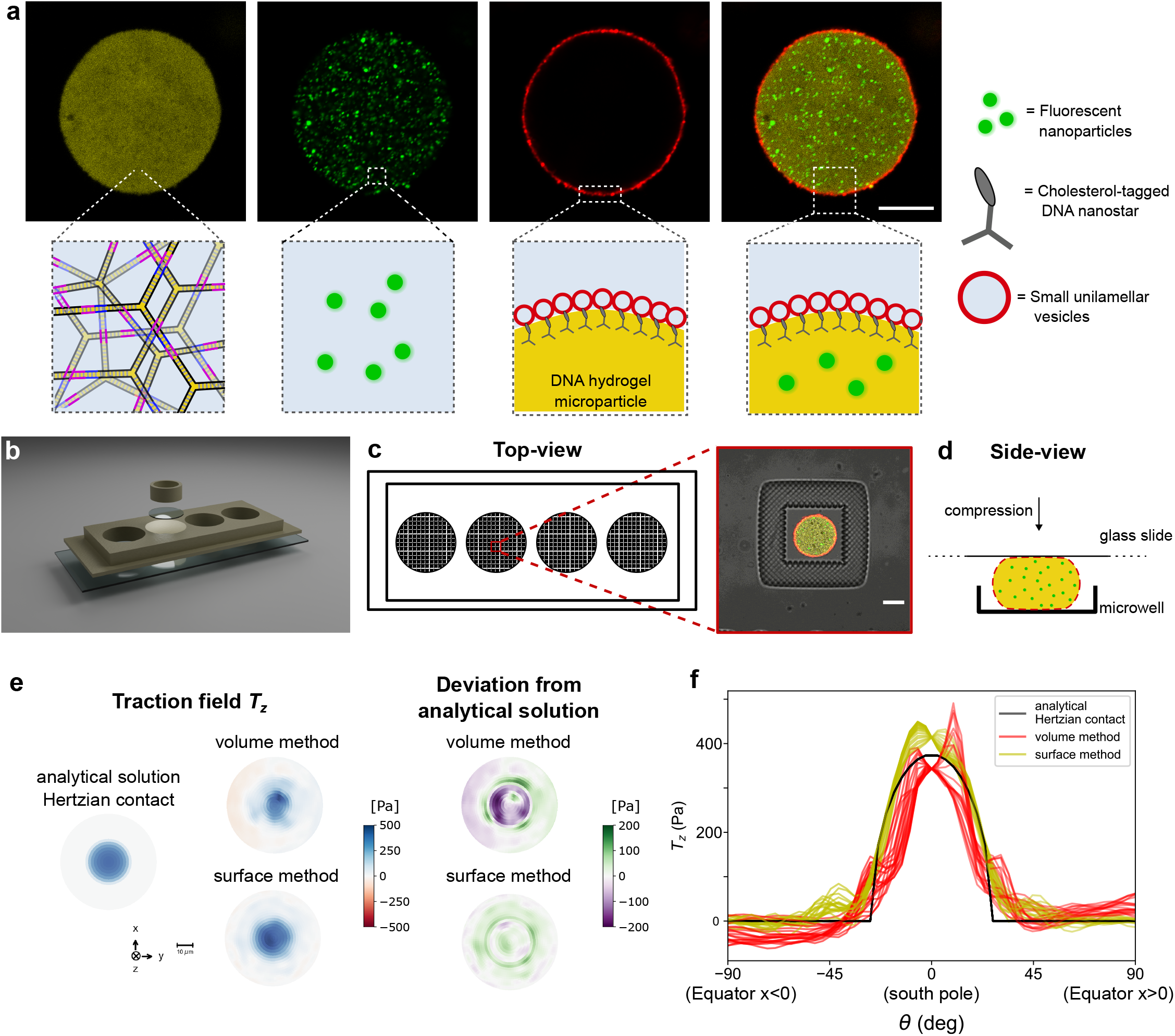
Design of experimental system and recovered traction profiles. (a) Confocal images (top row) and sketches (bottom row) illustrating the design and composition of DNA-HMPs engineered for this study. Confocal images show the DNA network composed of Cy3-labeled DNA nanostars (yellow, *λ*_ex_ = 561 nm) with embedded fluorescein-labeled nanoparticles (green, *λ*_ex_ = 488 nm) and coated with Atto-633-labeled SUVs (red, *λ*_ex_ = 640 nm). The final image shows a fluorescence overlay of all channels and components. The fluorescent nanoparticles thus label the volume; the DNA itself and the SUVs carry information about the surface shape, allowing for comparison of the methods on the same particle. Scale bar: 20 µm. (b) Exploded-view 3D rendering of the deformation setup as a custom 3D-printed 4-well holder (11 mm diameter wells), glued onto a glass slide and equipped with laser-printed PEGDA-based microwells of 40 µm diameter. Following DNA-HMP deposition into the microwells, the large wells were covered with a glass coverslide and the DNA-HMPs weighed down with a ring weight. (c) Sketch depicting the top-view of the observation chamber with the four large wells and printed microwells. The zoom image depicts a DNA-HMP deposited into one of the microwells. Scale bar: 20 µm. (d) Sketch depicting DNA-HMP deformation in a microwell by the deposited glass slide as a side-view. (e) Exemplary recovered traction profiles (bottom view on *T*_*z*_) for both methods from the same microparticle (assuming *E*_0_ = 1.1 kPa, *ν* = 0.4), and error/deviation from a Hertzian contact profile with a suitable contact radius (*a* = 0.45). (f) Superimposed cross-sections of traction profiles. Each line starts at the equator and finishes at the equator on the opposite side of the sphere.

To apply controlled forces, DNA-HMPs were trapped in microwells and compressed from above by a glass coverslide with increasing load facilitated by the addition of a 3D-printed weight (see Section 2.2.7). This way, it was possible to lower the applied force again after deformation by removing the weight (Fig.5(b-d)). As the glass slides are significantly stiffer (*E* ~ GPa) than the DNA-HMPs, the particle is expected to deform with a Hertzian contact profile as introduced above in Fig.2(a). The localization of microparticles in individual wells enabled us to take a reference picture of the uncompressed DNA-HMP for the volume method before or after compression.

We then applied the volume and surface method workflows established for the simulations to experimental data. Since our DNA-HMPs did not have a suitable indicator for traction-free regions for the surface method, this term does not contribute to the energy functional here, but should also not decrease performance significantly for the expected deformation levels.^29^ Fig.5(e, f)) show our results for one representative example. We note that the obtained traction profiles for the bottom side of the particles facing the microwell are similar in shape to the analytical Hertzian contact solution for a suitable contact radius (with *a* = 0.45). Evaluating the NMAE from the analytical solution across the bottom half of the sphere (z *<* 0) yields 35% for the surface method and 64% for the volume method, again showing a performance advantage of the surface method. However, the volume method appears to perform much closer to the optimum NMAE achieved in the simulations without noise (56% for *a* = 0.5) than that of the surface method (8% for *a* = 0.45, both see Fig.S15). We attribute this to the aforementioned robustness to noise of the volume method, where the NMAE is basically unaffected up to displacement noise values of 0.5%/px in the simulations. For the surface method, on the other hand, as shown in Fig.3, the NMAE strongly scales with noise, and assuming no other noise sources, the experimentally obtained NMAE for the given indentation (*a* = 0.45) suggests an effective surface roughness of approximately 1–2% for the surface method. Analyzing the surface roughness of the undeformed microparticle via the NRMS, we indeed find a value of ~ 2.6% for the bottom half of the sphere.

For the top side of the DNA-HMP, the traction reconstructions by both methods appear less accurate (Fig.S20), reflected in higher NMAE errors of 130% for the volume method and 82% for the surface method. We attribute the decrease in performance to a deterioration of resolution near the top of the surface, as evident by the recovered microparticle surface (Fig.S21); here the flat bottom part is cleanly resolved whereas the top part clearly shows visible roughness, which again directly translates into higher NMAE for the surface method. As the performance of the volume method appears to be affected as well, we assume a decrease in imaging quality (fluorescence signal compared to noise) towards the top, possibly caused by higher scattering, to be responsible for the performance decrease. This could be partially caused by the glass slide on the top, while it is pushed down by the weight, or by movements of the scanning microscope. This problem is common for z-stack imaging and has also been observed for polystyrene particles, inspiring the usage of deformable acrylamide-co-acrylic acid microparticles in the original surface method study.^29^ The particles used in that study were also more homogeneously spherical, with an NRMS of *<* 1%, which as in our case includes sphericity deviations caused by both imaging and variations in the particles themselves. The analyzed microscope data for this DNA-HMP is shown in Fig.S22. Beyond this, these results imply that with improved imaging and force control, reconstruction quality could be higher in future experimental setups. Further, because our force application occurred along the z-axis, along which resolution is typically lower in confocal microscopy, results might improve when considering a different axis for force application. Additional results are given in Fig.S23, showing again a spatial broadening of the traction profile for the volume method and similar problems on the top side of the microparticle compression setup.

Experimental implementation of the volume method proved challenging, mainly due to the simultaneous requirements of controlled force application, high-quality volumetric imaging, and stable reference image acquisition. Consequently, the number of microparticles which yielded reliable reconstructions was limited. For the surface method, deformed surface shapes could be reconstructed and traction profiles computed for more examined microparticles, again highlighting its broader applicability due to ease of experimental implementation. Additional microparticle profiles showing surface method evaluations are provided in Fig.S24. The traction fields mostly reflect the compression, but they also reveal that improved resolution and force application are necessary to cleanly resolve the expected Hertzian contact profile, particularly given that the surface method relies only on information about the deformed surface shape, which has to be captured at high quality. Despite sample size limitations, our experimental observations are consistent with the trends observed in our simulations: The volume method exhibits attenuation and spatial broadening in the reconstructed traction profile, while the surface method generally produces sharper reconstructions but remains more sensitive to imaging noise.

### 3.4. Comparison with nonlinear elasticity models

Some of the simulations performed represent strong compression scenarios ranging up to 30% in maximum relative indentation. The maximum relative displacement in the experiments are estimated to be up to 20%. This raises the question if the assumptions using linear elasticity theory (LET) still hold. LET is generally considered to be valid for strains up to *u*_*ij*_ ≲ 0.1. In Fig.6(a), we reconstruct the strains of the experimental data used in Fig.5. One sees that there is a typical compressive strain around 0.1 in the middle of the microparticle, which then reaches peak values of up to 0.2 closer to the surface, until going down to 0.0 at the two surfaces. We also note that the two sides are not equivalent, because the microparticles are placed on the bottom surface and loaded from the top.

**Figure 6.**
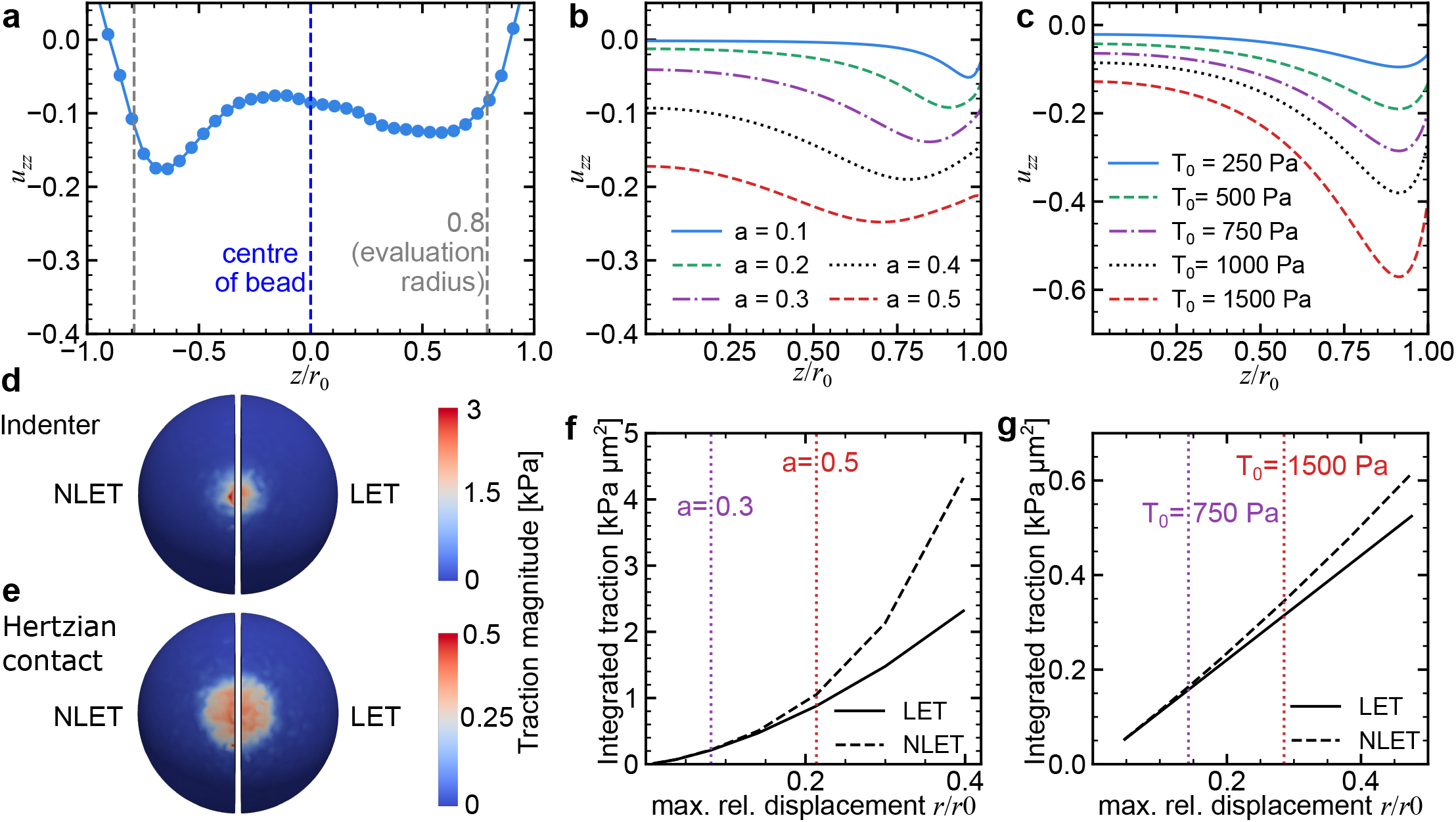
Limits of linear elasticity. (a) Experimentally obtained strain values of the full sized image along the z-axis. The evaluation radius is indicated by the dashed vertical lines. Same data as in Fig.5. (b) Analytical strain solution (*u*_*zz*_) for Hertzian contact scenario of different compression magnitudes, characterized by the contact radius *a*. Note that now we show only one half due to symmetry. The curves have the same shapes as the experimental data. (c) Analytical strain solution for the Gaussian indenter, compression characterized by maximum traction *T*. Here, the maximum observed strains are generally higher near the surface, but increase towards the center of the sphere, in line with the observations in Fig.4. (d) Gaussian indenter of high magnitude (maximum relative indentation 29 %), side-by-side comparison of traction profile recovered via FEM using non-linear elasticity theory (NLET, left) and linear elasticity theory (LET, right). Both yield similar results. (e) Side-by-side comparison of LET vs. NLET for Hertzian contact (maximum relative indentation 8 %), again showing similar results. (f) Integrated traction magnitude for Hertzian contact of FEM simulations, showing good agreement between LET and NLET up to contact radius *a* = 0.5. (g) Integrated traction magnitude for indenter scenario, showing good agreement even at high compression.

The experimentally measured curve agrees well with the simulation results for the Hertzian contact shown in Fig.6(b), which by construction are symmetric in bottom and top, thus we only show one side. For the indenter scenario, the strains observed in the simulations range even higher, up to 0.6 for the maximum indentation level (Fig.6(c), T = 1500 Pa).

To test if the assumption of linear elasticity leads to errors in the traction estimation, we performed finite element method (FEM) simulations comparing LET and NLET, the latter implemented by a hyperelastic (neo-Hookean) model. As shown in Fig.6(d-g), LET and NLET yield similar results up to a maximal relative displacement of 0.2, which is the level reached in our experiments. The experimentally estimated contact radius (*a* = 0.45) is just below the limit of when the models start to diverge for the Hertzian contact (Fig.6(f)), showing that even though the strain magnitude exceeds 0.1, the assumptions of LET still seem to hold. For the indenter scenario, the agreement seems to persist even to higher levels (Fig.6(g)).

## CONCLUSIONS

In this work, we performed a systematic comparison of the two most common strategies used in MP-TFM: the volume method and the surface method. Using simulated traction profiles representing three biologically relevant loading scenarios, we found that considering similar noise levels, the surface method consistently reconstructs traction fields with substantially lower errors than the volume method. Although it might appear difficult to compare two methods that are very different in nature, we note that both can be considered to be regularized and that both have been optimized for MP-TFM.

Analysis of the displacement fields revealed that this performance difference originates primarily from surface artifacts inherent to the digital volume correlation algorithm used in the volume method. While the recovered displacement field remains accurate throughout most of the particle interior, tracking accuracy deteriorates close to the particle surface due to the lack of fiducial markers outside the particle. Because traction reconstruction requires spatial derivatives of the displacement field, these small deviations become strongly amplified near the boundary. Consequently, traction fields must be evaluated at radii further inside the particle, which leads to a loss of information near the deformed surface where the forces are applied.

We further provide quantitative error estimates across a range of noise levels and loading strengths for the volume method, the surface method, and a simplified “truncated” surface approach that omits displacement optimization, offering faster computation at the expense of accuracy. These results establish practical benchmarks for experimental MP-TFM studies. Our analysis shows that the volume and surface methods exhibit different sensitivities to measurement noise. The surface method achieves significantly lower reconstruction errors under low-noise conditions but shows a strong dependence on surface roughness. In contrast, the volume method is substantially more robust to displacement noise of the embedded nanoparticles because the correlation algorithm averages information from many markers within the particle volume. As a result, the performance gap between the two methods decreases at high surface roughness noise levels, making the surface method more sensitive to imaging conditions and to the sphericity of the used microparticles. For future applications, it is therefore advisable to improve the performance of the surface method by establishing more accurate estimates of the reference configuration. It is left to future work to decide which method performs best in the context of complex and time-dependent processes in real tissues, for which confounding effects like scattering have to be taken into account. Here it might be interesting to develop new pipelines, like an inverse regularized method for the volume method, especially in light of its success in classical 2D-TFM.

To experimentally validate these findings, we implemented both approaches using DNA-HMPs equipped with fluorescent surface labels and embedded nanoparticles. Compression of the particles in a custom microwell setup produced traction profiles consistent with a Hertzian contact model, and allowed for reconstruction by both methods. In agreement with the simulations, the surface method yielded lower reconstruction errors, although the difference between the two approaches was reduced under experimental conditions due to imaging noise and the lower noise sensitivity of the volume method. The lower noise sensitivity partially stems from the necessity of the reference image for the volume method, but generally both methods profit from high-quality microscope images. Still, because the surface method does not need a reference image, it proved easier to apply reliably, as it is independent of image alignment or unintended nanoparticle motion, which could occur due to microparticle rotation.

Beyond the method comparison, our results demonstrate that DNA-HMPs provide a versatile platform for MP-TFM experiments. Their tunable mechanical properties and compatibility with multiple fluorescent labels enable simultaneous implementation of both traction reconstruction strategies. The lipid SUV coating enables further biological functionalization, e.g., with membrane proteins like E-cadherins and thus the creation of more cell-like interfaces, which may facilitate future studies of cellular force generation. As our experimental analysis was limited to the Hertzian contact scenario, more experimental work is needed to verify our simulation predictions for the ring and indenter scenario, even though it might be difficult to obtain a ground truth in these cases. Another interesting avenue of investigation is the use of viscoelastic materials.

Overall, our results suggest that even though the volume method in principle relies on fewer assumptions and uses more information, namely the reference image and the full displacement field, the surface method is generally the preferred approach for MP-TFM due to its higher reconstruction accuracy and simpler experimental implementation. However, the volume method may remain advantageous in situations where surface imaging is compromised, such as near optical interfaces or in strongly scattering environments. Our findings provide practical guidance for selecting and interpreting traction reconstruction methods in future MP-TFM studies.

## AUTHOR CONTRIBUTIONS

S.B., B.K.K., T.W., C.M., and T.A. performed the research. S.B., B.K.K., and C.M. wrote the computer code and performed the data analysis. B.K.K., T.W., and T.A. performed the experiments. S.B. wrote the original draft of the manuscript, supported by T.W. K.G. and U.S.S. conceptualized and supervised the research. All authors reviewed the manuscript.

## CONFLICTS OF INTEREST

There are no conflicts to declare.

## DATA AVAILABILITY

We provide a Python script and necessary files to run simulations of a Hertzian contact scenario with evaluations by the volume method and the surface method, a JupyterLab notebook for surface method reconstructions of deformed DNA-HMPs, and a JupyterLab notebook supplemented by exemplary displacement data and results for the FEM comparison of linear elasticity and hyperelasticity, on GitHub (https://github.com/brewburgr/MPTFM). The experimental data is available at https://doi.org/10.11588/DATA/KWOD5A.

## ACKNOWLEDGEMENTS

The authors thank Johannes Blumberg and Leon Lettermann for helpful discussions. The authors thank Dr. Sadaf Pashapour and the Microfabrication and Microfluidic Core Facility (µFluCF) at the Institute of Molecular Systems Engineering and Advanced Materials (IMSEAM) funded partly by the Health + Life Science Alliance Heidelberg Mannheim. The Health + Life Science Alliance provided state funds approved by the State Parliament of Baden-Württemberg. S.B. acknowledges funding through EP/X038009/1 Horizon Europe UKRI Underwrite MSCA ‘DYNAMO’ and thanks Prof. Ulrich F. Keyser for support. T.W. thanks the German National Academic Foundation. K.G. and U.S.S. acknowledge funding from the Deutsche Forschungsgemeinschaft (DFG, German Research Foundation) under Germany’s Excellence Strategy via the Excellence Cluster 3D Matter Made to Order (EXC-2082/1-390761711 and EXC-2082/2-390761711), SynthImmune (EXC-3018/1 – 533587280) and the Collaborative Research Center Membrane remodelling (SFB-1638/1–511488495). K.G. acknowledges the ERC Starting Grant ENSYNC (101076997).

## SUPPLEMENTARY INFORMATION (SI)

**Figure S1.**
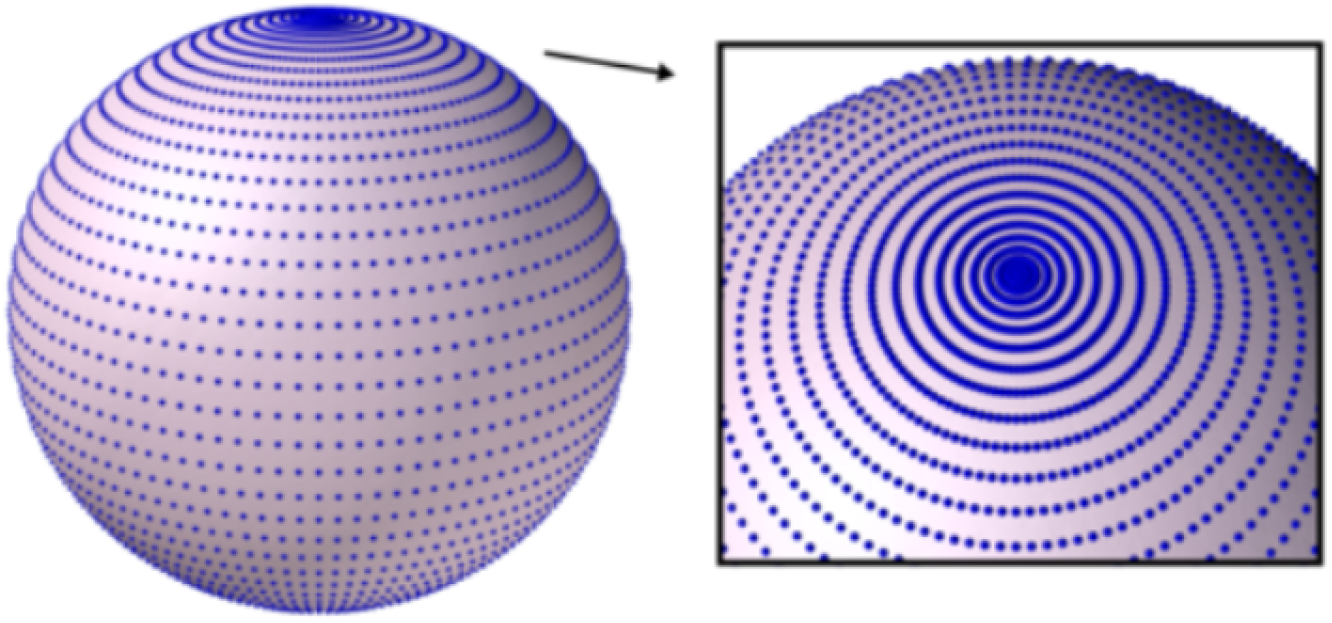
Visualization of a Gauss Legendre quadrature (GLQ) mesh.^48^ This mesh is used for the minimization and traction evaluation in the surface method, and it allows for an efficient workflow together with the spherical harmonics (SH) approach.^29,36^

**Figure S2.**
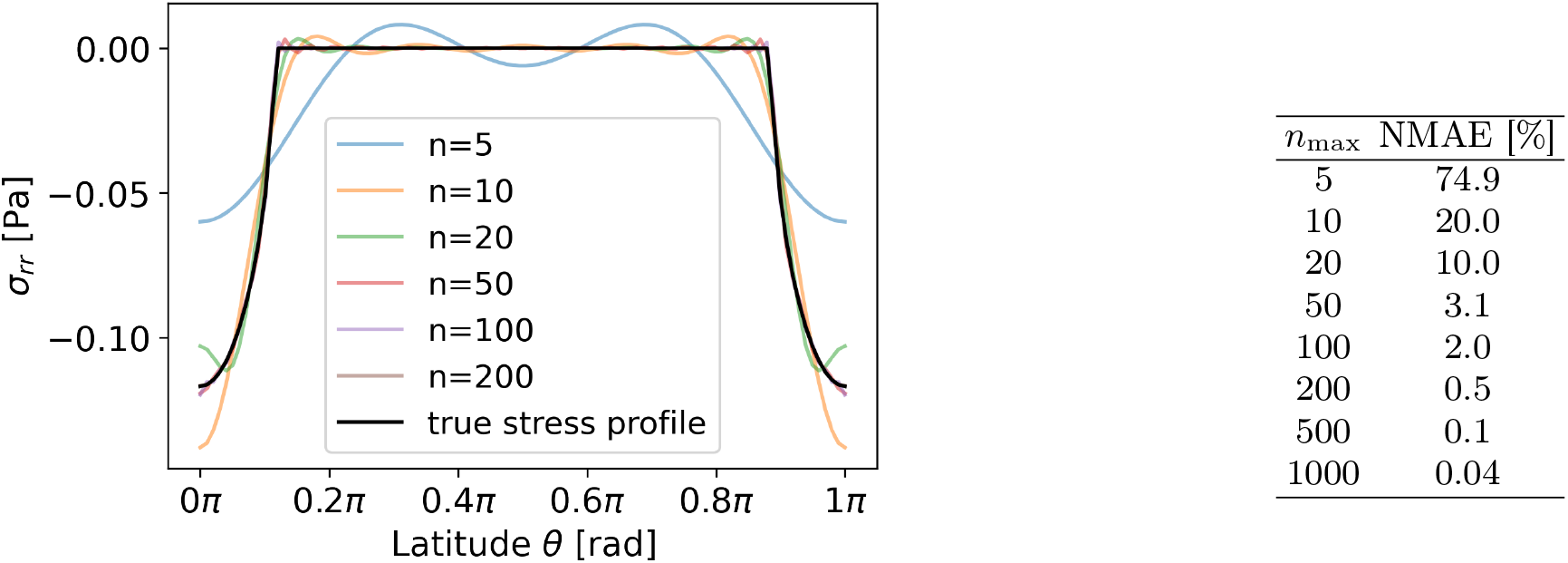
Decomposition accuracy test. (a): True and reconstructed stress profiles *σ*_*rr*_ for a Hertzian contact scenario. The cutoff *n* = *n*_max_ is the maximum sum order used in the decomposition of the analytical solution (see Eq.(10)^44^). (b): Normalized mean average error (NMAE) for different *n*_max_. For *n* ≥ 50, the deviations are sufficiently small in comparison to the errors expected by the methods themselves. We choose a cutoff *n*_max_ = 50, which provides an acceptable 3% deviation for the Hertzian contact profile at reasonable runtimes for the simulation of the experimental data (𝒪 (1 min)). As the Hertzian contact profile has a sharp edge, which requires more higher order terms to be resolved accurately, we can expect the results for smoother profiles, e.g., the Gaussian indenter or ring, to be even more accurate.

**Figure S3.**
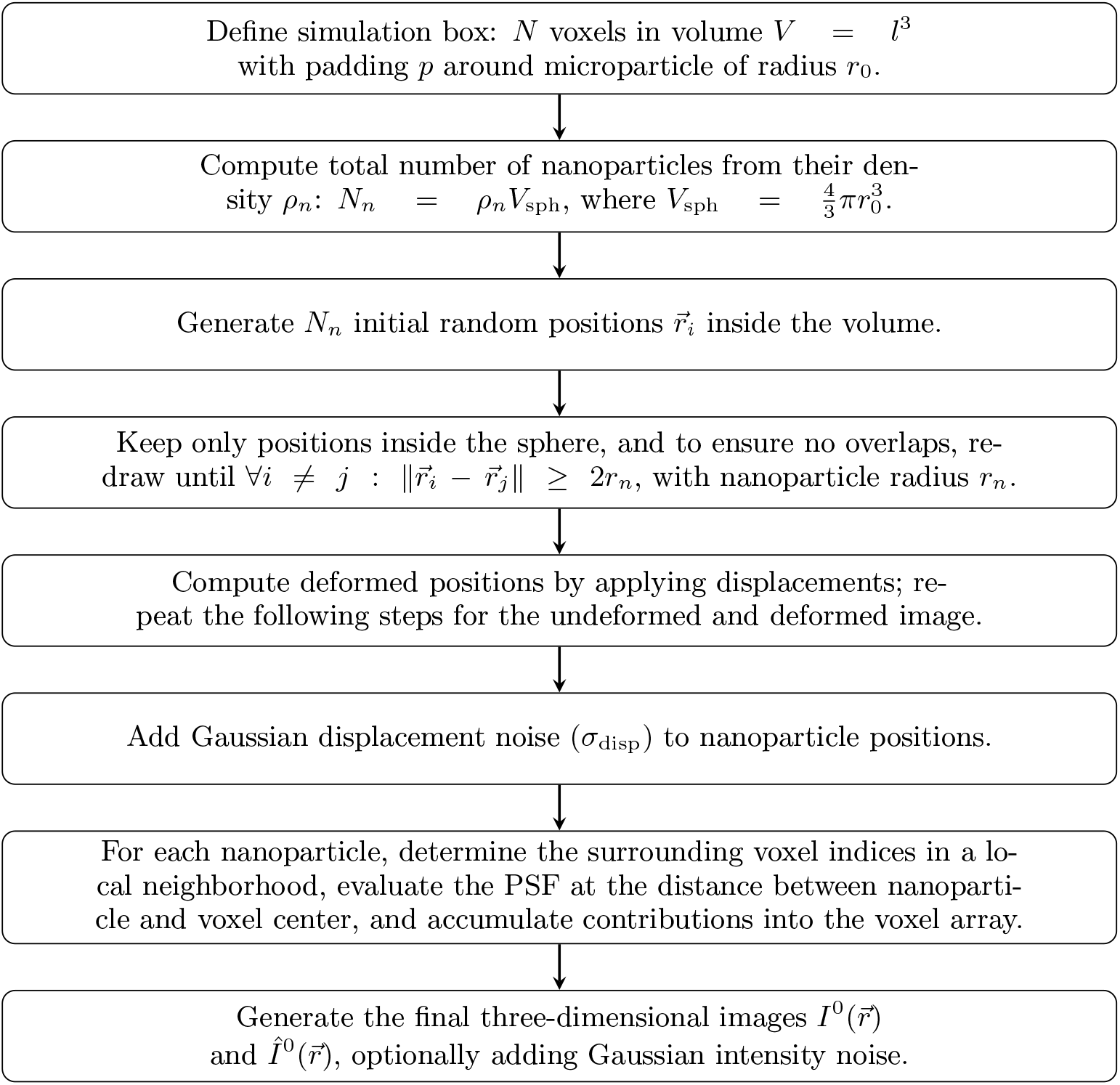
Workflow for generating synthetic reference and deformed three-dimensional nanoparticle images for the volume method. Steps include volume definition, nanoparticle placement with minimum distance and inside the microparticle, displacement application, Gaussian displacement noise, and PSF-based intensity computation. The redrawing in step 4 is computationally expensive as the distance matrix to all other points needs to be calculated, and could be skipped, allowing for nanoparticle overlap. For standard parameters, given in Table S1, the runtime is in the range of seconds.

**Figure S4.**
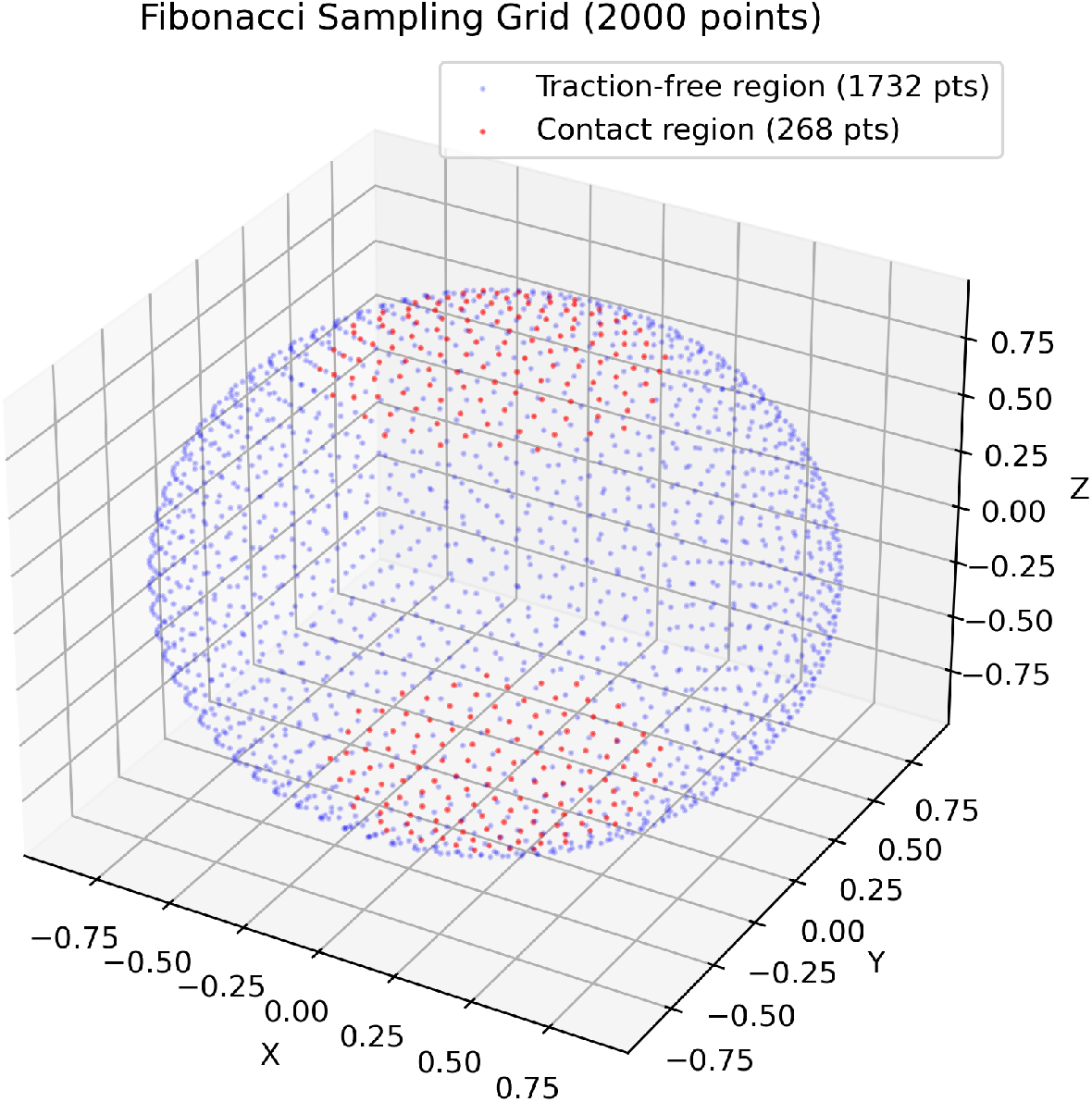
Visualization of a (deformed) Fibonacci grid, exemplary synthetic data for the surface method workflow. The set of points describing the microparticle surface is used to simulate a Hertzian contact (*a* = 0.5) with *N* = 2000 Fibonacci samples (here normalized to a unit sphere), and shown is the polar area of force application (red) as well as the region known to be traction-free (by the prescribed traction profile, see Note S4) used for the functional minimization (blue). Sampling on a Fibonacci grid leads to almost even spacing between the points, and the oversampling at the poles that a Gauss-Legendre quadrature (GLQ) would introduce is prevented. This oversampling would be advantageous for Hertzian contact and indenter profiles with force application at the poles, but disadvantageous for ring profiles, where force is applied in the equatorial region.

**Figure S5.**
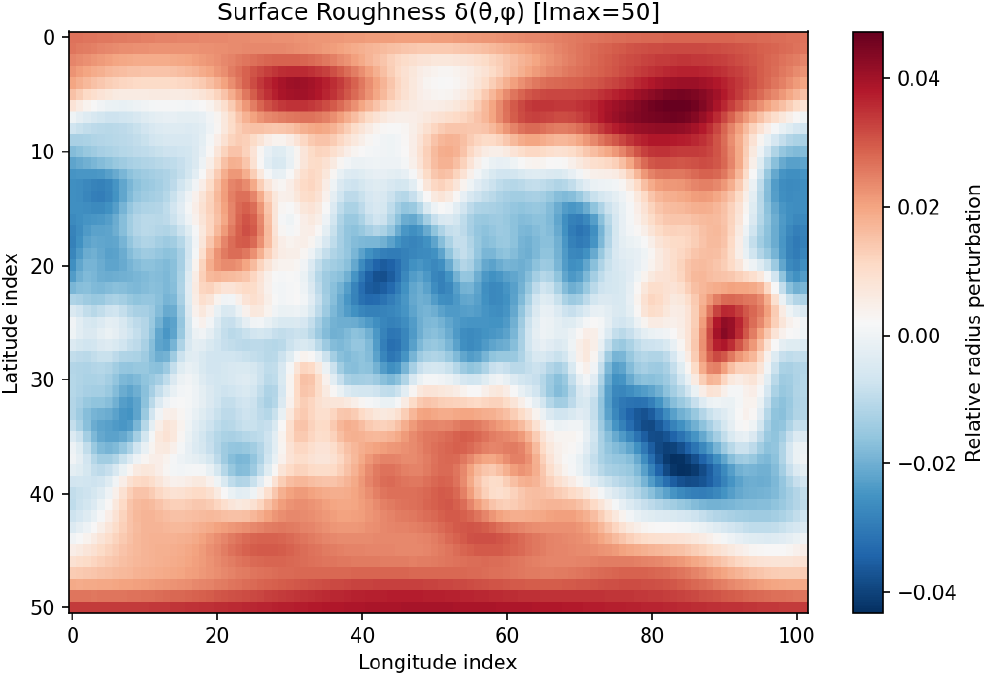
Surface roughness noise generation for surface method simulations. Visualization of an exemplary surface roughness distribution *δ*(*θ, φ*), implementing radial perturbations with a spectral approach, here with a prescribed root-mean-square (rms) of 0.02 (relative to the reference radius *r*_0_). This approach is used to simulate noise for the surface method as surface roughness, which can be quantified compactly by the rms. Indices cover the sphere but do not reflect degree or radian units.

**Figure S6.**
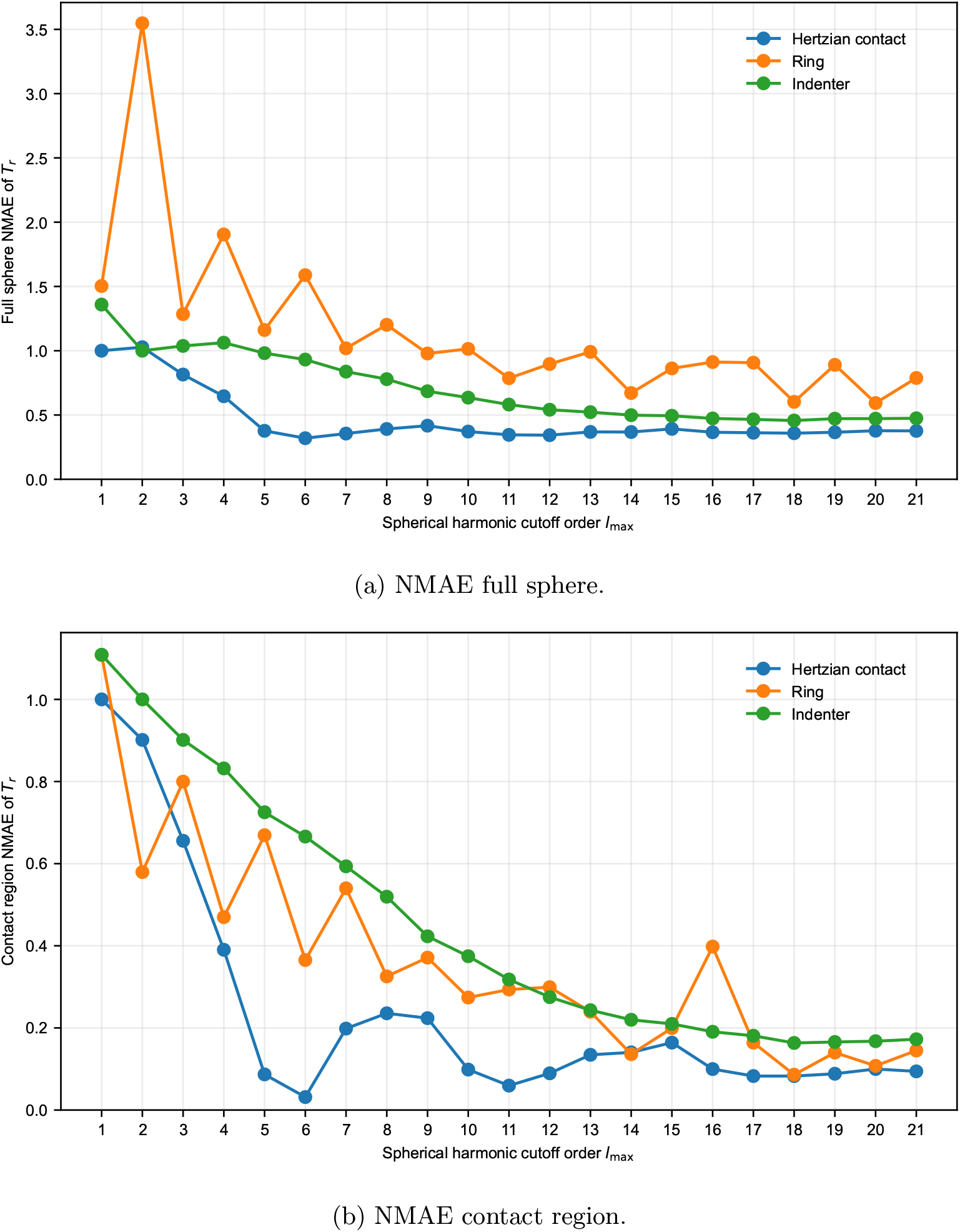
Surface method spatial resolution analysis. We evaluated traction reconstructions with respect to the spherical harmonic cutoff order *l*_max_ for the minimization, evaluated either over the full sphere (a) or within the contact region only (b). The normalized mean absolute error (NMAE) of the reconstructed traction field component *T*_*r*_ is shown as a function of harmonic cutoff order for the examined deformation scenarios (Hertzian contact, ring, indenter). Reconstruction accuracy improves overall with increasing harmonic order, with errors decreasing rapidly at low *l*_max_ before approaching a plateau around *l*_max_ ≈ 18. Beyond *l*_max_ *>* 20, only marginal accuracy improvements are observed. At the same time, computational cost increases, scaling approximately with 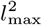. These findings motivate the use of *l*_max_ = 20 as the standard reconstruction parameter throughout this work, providing a suitable balance between reconstruction accuracy and computational efficiency. Less localized traction profiles, such as Hertzian contact, converge at lower harmonic orders (*l*_max_ ≈ 15), whereas highly localized traction distributions, such as the indenter geometry, require higher cutoffs to resolve sharper spatial features. More localized experimental force distributions may require higher harmonic orders together with correspondingly high-resolution surface reconstructions to avoid overfitting and aliasing (see also Fig.S14). Characteristic Gibbs-like oscillations are observed near the ring equatorial edge at low harmonic orders and diminish with increasing *l*_max_, illustrating why localized traction profiles require higher harmonic resolution than smoother force distributions.

**Figure S7.**
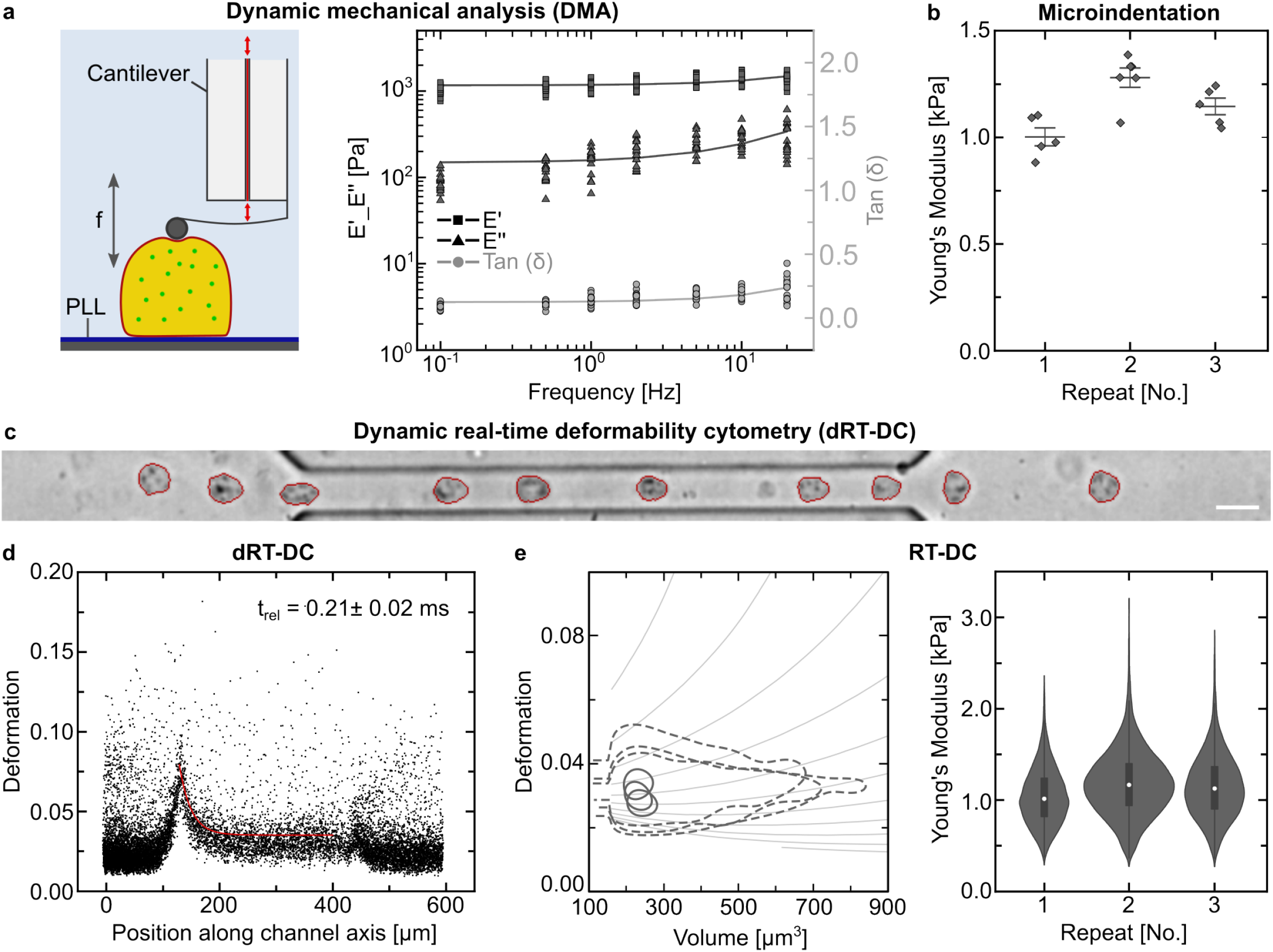
Mechanical characterization of DNA-HMPs for MP-TFM. (a) Schematic outlining DNA-HMP analysis via dynamic mechanical analysis (DMA) with DNA-HMPs attached to a glass substrate using electrostatic interaction following PLL-coating of the glass slide. Graph depicting DMA data of the DNA-HMPs. The storage modulus *E*^*′*^ (squares) and the loss modulus *E*^*′′*^ (triangles) are plotted on a logarithmic scale, tan(*δ*) (*E*^*′′*^*/E*^*′*^, filled circles) is plotted on a linear scale. (b) Analysis of Young’s modulus of the DNA-HMPs as extracted from Hertzian-fits to microindentation curves on the DNA-HMPs. The data is depicted as individual data points for each of the triplicate measurements also showing the mean ± standard deviation. (c) Composite image of the DNA-HMPs in the flow channel during dynamic real-time deformability cytometry (dRT-DC). The DNA-HMPs are initially spherical prior to entering the flow channel and get deformed strongly upon insertion into the channel. Traversing the channel, the DNA-HMPs relax into a steady-state deformation and return to their spherical shape after leaving the channel. Scale bar: 20 µm. (d) Deformation over position along channel axis scatter plot of the DNA-HMPs during dRT-DC. Each point corresponds to an individual DNA-HMP (n = 13000). The relaxation time t_rel_ of the DNA-HMPs was extracted from the exponential fit of the relaxation scatter plot (red curve) as previously described.^38^ (e) Analysis of DNA-HMPs via real-time deformability cytometry (RT-DC). Contour plot depicting DNA-HMP steady-state deformation over particle volume for triplicate measurements showing the 50^th^ percentile (dashed line) and 95^th^ percentile (solid line). Isoelasticity lines derived from numerical simulations are shown additionally, indicating stiffness changes where a steeper slope corresponds to softer particles. The data depict strong overlap and thus good reproducibility between replicates. Young’s modulus of the DNA-HMPs as derived from RT-DC steady-state measurements. Plot depicting the measured Young’s modulus of n = 3 replicates of the DNA-HMPs (n_1_ = 8913, n_2_ = 16021, n_3_ = 10635). The data is presented as violin plots showing the median value (white dot) of each measurement with boxplots encompassing the 25 - 75 % percentiles and a whisker length of 1.5 IQR. Mechanical analysis of the DNA-HMPs was conducted via dynamic mechanical analysis (DMA) by microindentation as well as real-time deformability cytometry (RT-DC) as outlined in a previous publication.^38^ The DNA-HMPs were adhered to a glass substrate following electrostatic interaction with poly-l-lysine (PLL, MW = 150 kDa - 300 kDa) and measured with increasing indentation frequency using a spherical cantilever tip (Fig.S7a). DMA revealed largely frequency-independent storage moduli E’ for the DNA-HMPs used in this study. Further, *E*^*′′*^ remained below *E*^*′*^ for all tested frequencies with the loss tangent tan(*δ*) staying below 0.3 even at the highest frequency tested (20 Hz). This denotes the DNA-HMPs used here as behaving predominantly elastic. The mean Young’s modulus *E* of the DNA-HMPs across triplicate measurements was further measured at (1142 ±169) Pa (Fig.S7b). Additionally, we performed dynamic real-time deformability cytometry (dRT-DC) at 0.04 µL*/*s total flow rate on the DNA-HMPs. As shown in Fig.S7c/d, deformation of the particles peaked upon channel entry, showing quick relaxation of the DNA-HMPs into a bullet-shaped steady-state deformation around 0.035 (Fig.S7c/d/e). An exponential fit to the slope of DNA-HMP relaxation scatter plot following initial deformation then revealed a characteristic response time *τ* of (0.21 ± 0.02) ms. Further, extraction of the Young’s modulus *E* of the DNA-HMPs as measured via RT-DC at 0.04 µL*/*s total flow rate showed a mean value of (1117 ±720) Pa across the three triplicates and thus good agreement with the microindentation data (Fig.S7e). Calculation of the apparent viscosity of the DNA-HMPs used here following *τ* = *η/E*^38^ showed a low viscosity of 0.245 Pa·s, underlining the predominantly-elastic behavior of the DNA-HMPs. The marked increase in stiffness of the DNA-HMPs used in this study compared to their non-modified counterparts reported earlier^38^ can be explained by the addition of the fluorescent nanoparticles and SUV-coating; both of which can be expected to result in a stiffer network, yet do not seem to impair overall network flexibility.

**Figure S8.**
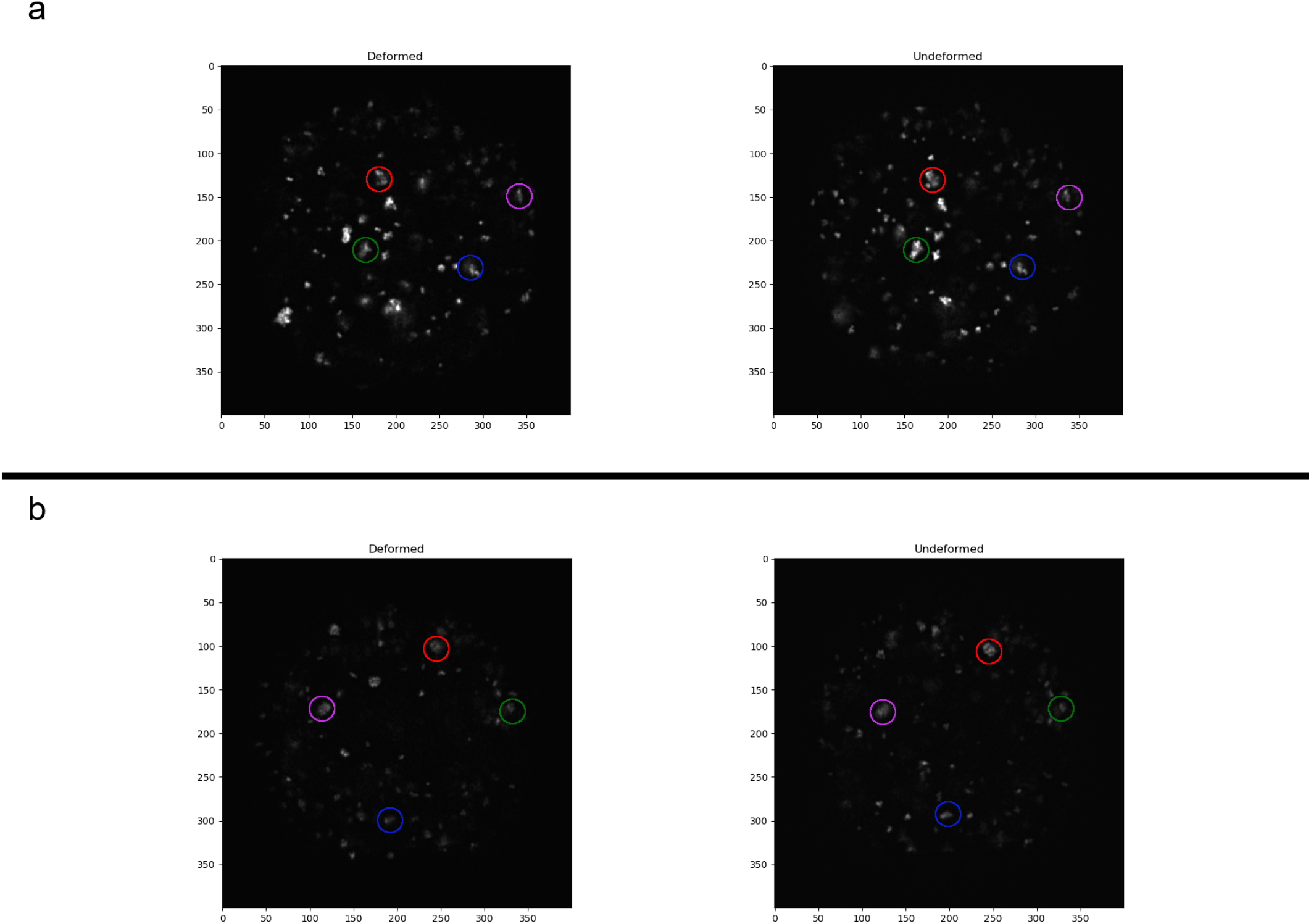
Visualization of preprocessed nanoparticle microscope images for the volume method at two different representative heights in the z-stack, consisting of 157 slices total. Left-hand side: Deformed configuration. Right-hand side: Undeformed (reference) configuration. (a) z = 65. (b) z = 105. In these aligned images, it is noticeable how several distinct nanoparticles moved from the undeformed to the deformed configuration, some of which are marked by colored circles. Units are voxels. The microparticle is the same one as in Fig.S9 and Fig.S10, and was analyzed in Fig.5(e, f). Preprocessing for the volume method. Here, the microscope images of the nanoparticles inside the DNA-HMPs in reference and deformed states are first preprocessed to allow for accurate correlation tracking. A Python pipeline has been developed, which first converts the nanoparticle images from .czi to .mat files, suitable for the correlation tracking algorithm (FIDVC), which runs in Matlab.^40^ Special care must be taken with the coordinate ordering, as microscope images are typically saved as arrays with coordinate order (z, y, x), while Python programs usually employ (x, y, z), and Matlab uses (y, x, z). The workflow starts by loading in two files (reference and deformed), and optionally padding them symmetrically with zero intensity stacks along the z-dimension. This can be done because typically the deformed microparticle image will contain fewer z-stacks than the undeformed one because it will be compressed along the z-axis, even though during microscopy, a few extra z-stacks are captured above and below the microparticle to ensure it is fully imaged. Usually the smaller value of the two z-dimensions between the two images is chosen as the objective z-dimension after cropping, but especially for larger deformations, the padding step is important as to not cut off the reference microparticle with fewer z-stacks later, which would lose information. In these cases, adding 10-20 voxels in z as a buffer could improve results for the present work. Then, a defined microparticle intensity threshold is used for a boolean mask of voxels assumed to be part of the physical microparticle, approximating its shape. Next, the intensity-weighted center of mass of both microparticles is computed in three dimensions, employing the mask. Bounding boxes are calculated by generating start and end indices centered on the center of mass, but clamped to stay inside the volume. The objective size in the x- and y-dimension is defined as 400 voxels each for the present work, which is typically sufficient to capture the entire microparticle. Because the center of masses might be different, the resulting windows can slightly differ. To ensure the dimensions of the resulting images match exactly, the crop size is defined as the minimum overlapping size in each dimension, reducing both bounding boxes to the same shapes. Finally, both volumes are cropped by the computed and clamped indices. An appropriate threshold is applied to suppress background noise while not losing information, and images are saved as .mat files for correlation tracking. Visualizations of the images throughout the z-stacks as well as intensity histograms are plotted throughout to check intermediary and final results, and tune parameters. It was found that the tracking algorithm and recovered displacements depend sensitively on properly aligned images. The two z-stacks of aligned images that are visualized show that singular nanoparticles can already be tracked by eye, and a meaningful displacement field for the volume method could be extracted by the FIDVC.

**Figure S9.**
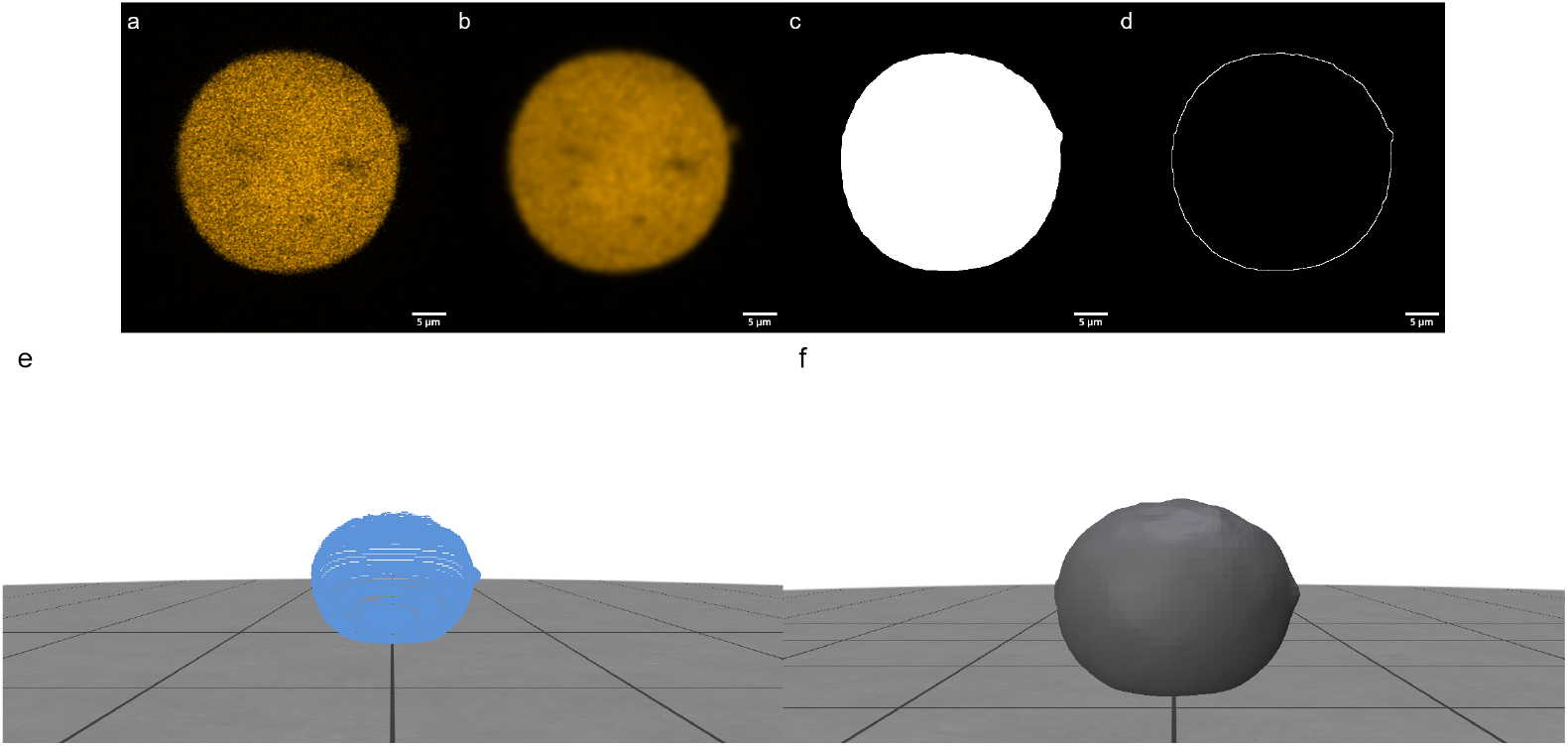
Preprocessing for the surface method, which requires the surface shape of a deformed microparticle. (a) Central z-stack of the original microscope image of the DNA network structure (Cy3). (b) A Gaussian blur is applied. (c) A threshold is applied by Otsu’s method. (d) After filling holes, an outline can be computed. (e) Visualization of the resulting set of points describing the microparticle surface. (f) Reconstruction of the surface shape by Poisson’s method after preprocessing and calculating normals. (a-d) are performed in Fiji^46^ (scale bars 5 µm). (e, f) are performed in GeoV.^47^ The set of points can be saved as a .ply file. Analysis results for this microparticle are shown in Fig.5(e, f). Preprocessing for the surface method. Here, in order to analyze the DNA-HMPs, the data was preprocessed to yield a reconstructed DNA-HMP surface, using the fluorescent Cy3 labels in the DNA network to recover a set of points describing the microparticle surface. First, the acquired z-stacks were opened in ImageJ 1.54f (NIH^46^), filtered using a Gaussian blur filter (*σ* = 4.0), and threshold-adjusted using Otsu’s method with parameters suitable for all z-stacks to yield binarized images. After that, the outline of the resulting binarized image stack was created after filling holes using Fiji’s inbuilt functions, and saved as a .tif file. Surface reconstruction was then conducted using the software GeoV.^47^ Within the software, the resulting set of 40,000 points was simplified to 0.4% world units and the data pre-cleaned to deplete outliers (0.5 probability). Surface normals were calculated with 2 smooth iterations, and the surface reconstructed using the screened Poisson method with a reconstruction depth of 5. All other parameters were kept as default. Parameters could be tuned in Fiji and GeoV, to yield smooth surface reconstructions while avoiding holes; though care must be taken not to falsify the observed surface shape. The resulting surface was then saved as a .ply file for deformation analysis with the surface method. The process is visualized for an example microparticle in Fig.S9, which also reflects differences in signal quality at the top and bottom of the microparticle.

**Figure S10.**
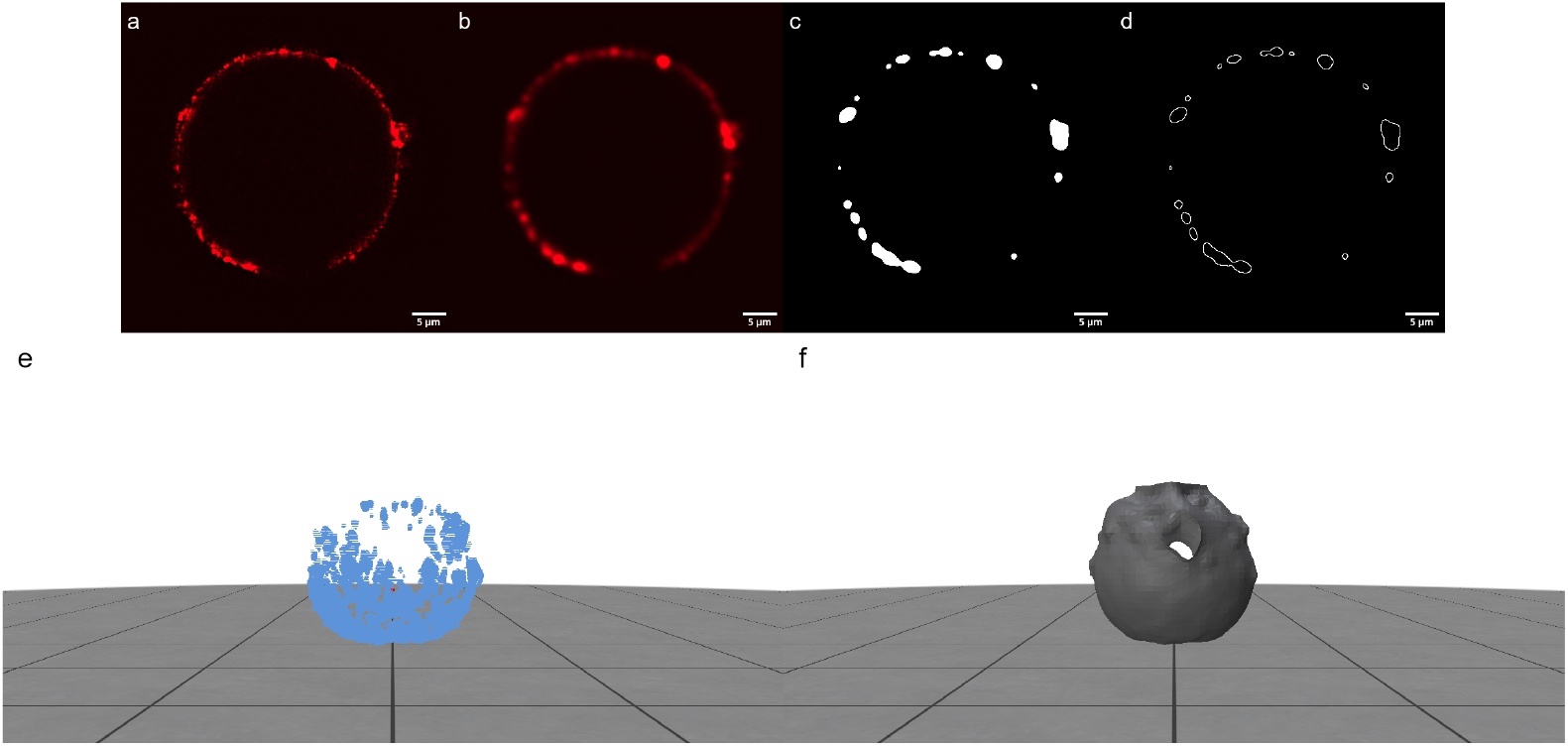
Attempted preprocessing of the surface label by the lipid SUV coating for the same microparticle as in Fig.S9. (a) Central z-stack of the original microscope image, showing the microparticle surface with small irregularities at high resolution. (b) A Gaussian blur is applied. (c) A threshold is applied by Otsu’s method. (d) After filling holes, an outline can be computed, but the surface is not clearly resolved. (e) Visualization of the resulting set of points describing the microparticle surface, containing holes and missing points at the top due to the optical setup, which also led to problems defining consistent thresholds. (f) Reconstruction of the surface shape by Poisson’s method from the set of points after preprocessing and calculating normals. (a-d) are performed in Fiji^46^ (scale bars 5 µm). (e, f) are performed in GeoV.^47^ The irregular surface containing holes is problematic, and it is not simple to smooth it out over all z-stacks. It does not reflect the deformed microparticle shape and cannot be used for the surface method. There have also been attempts to analyze the SUV coating (red), which was meant to serve as a surface indicator and could be useful in further biological setups, similar to the antibody coating for the traction-free region in the original publication.^29^ However, the surface could not be smoothly reconstructed this way. The main issue seems to lie in the fact that the resolution of the employed microscope is too high, leading to small irregularities and holes in the surface reconstruction, which were difficult to smooth out consistently over all z-stacks; moreover, the optical setup and photobleaching affected the reconstruction. The process is visualized for an example microparticle in Fig.S10. However, as Fig.S9 shows, reconstructions are possible using the Cy3 labels.

**Figure S11.**
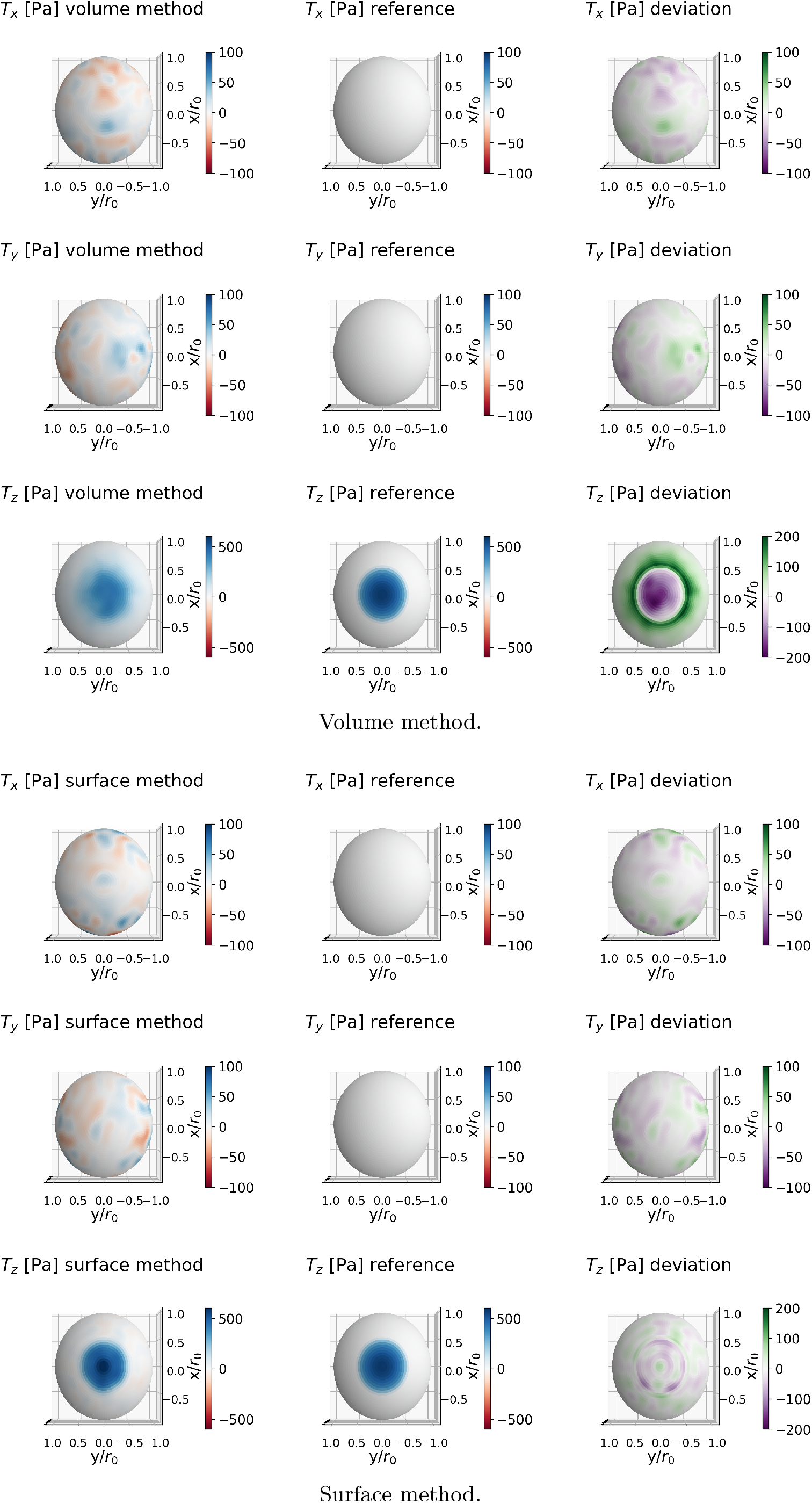
Full simulation results of all three traction components for the Hertzian contact scenario.

**Figure S12.**
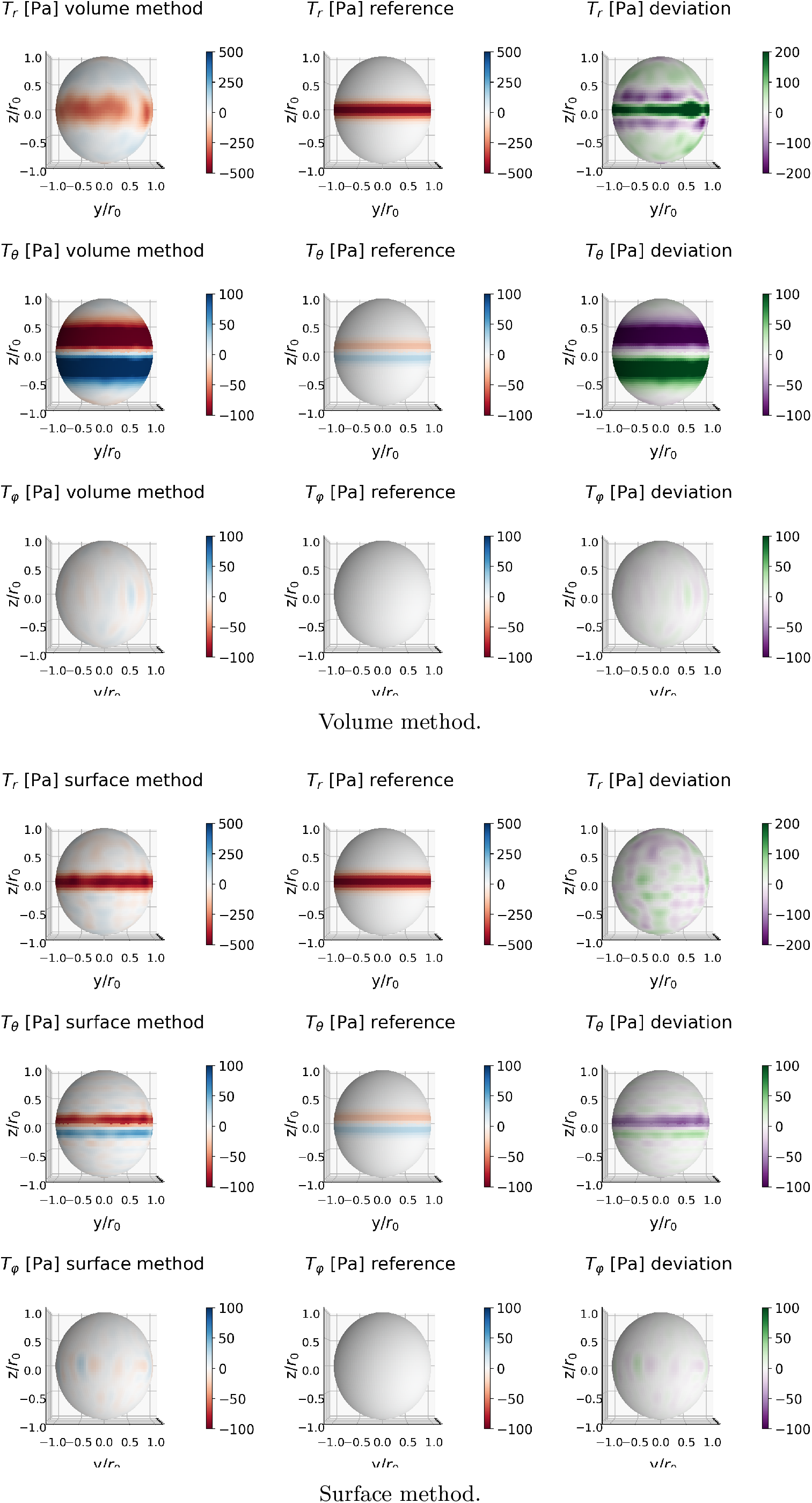
Full simulation results of all three traction components for the ring scenario.

**Figure S13.**
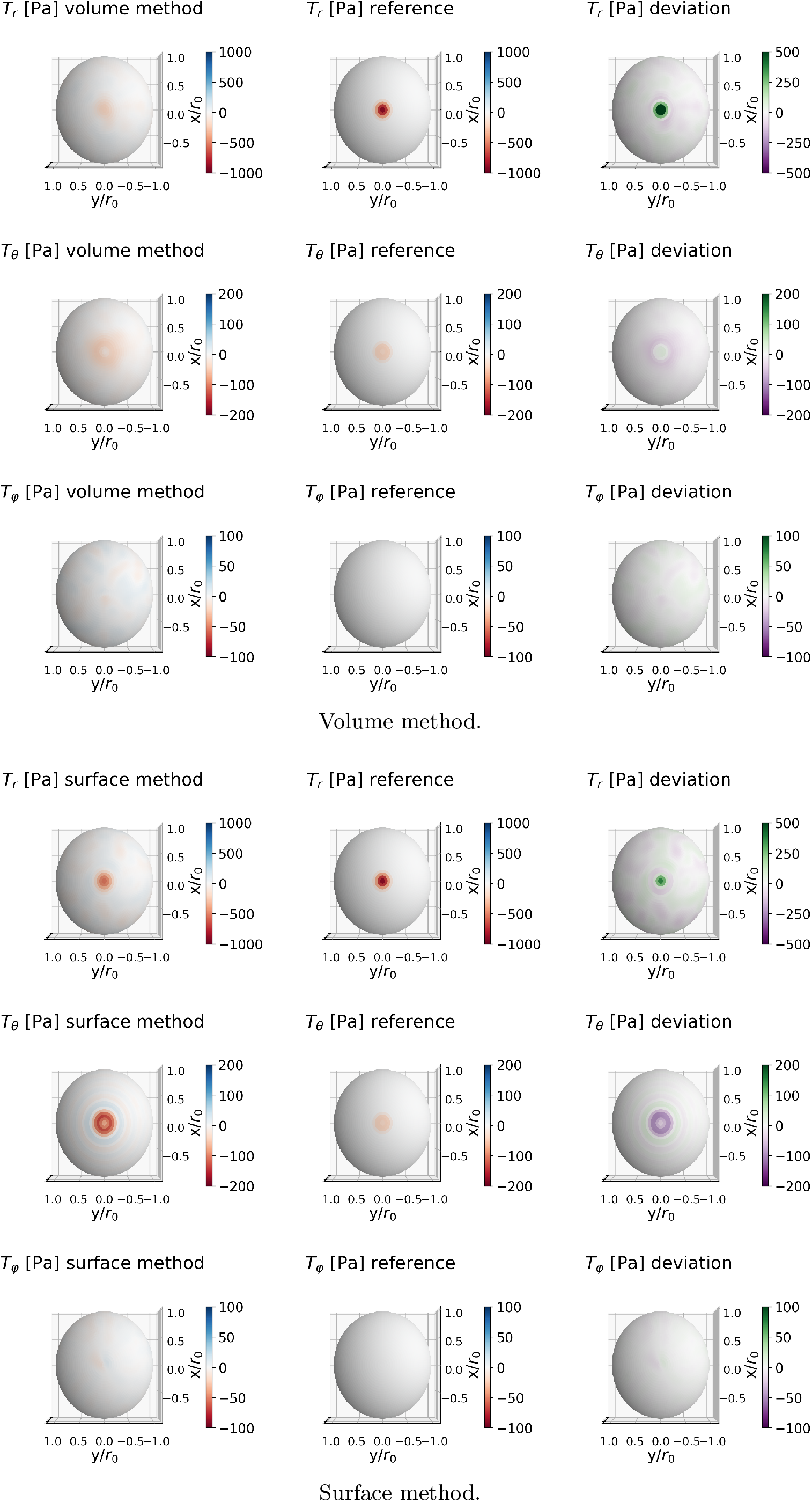
Full simulation results of all three traction components for the indenter scenario.

**Figure S14.**
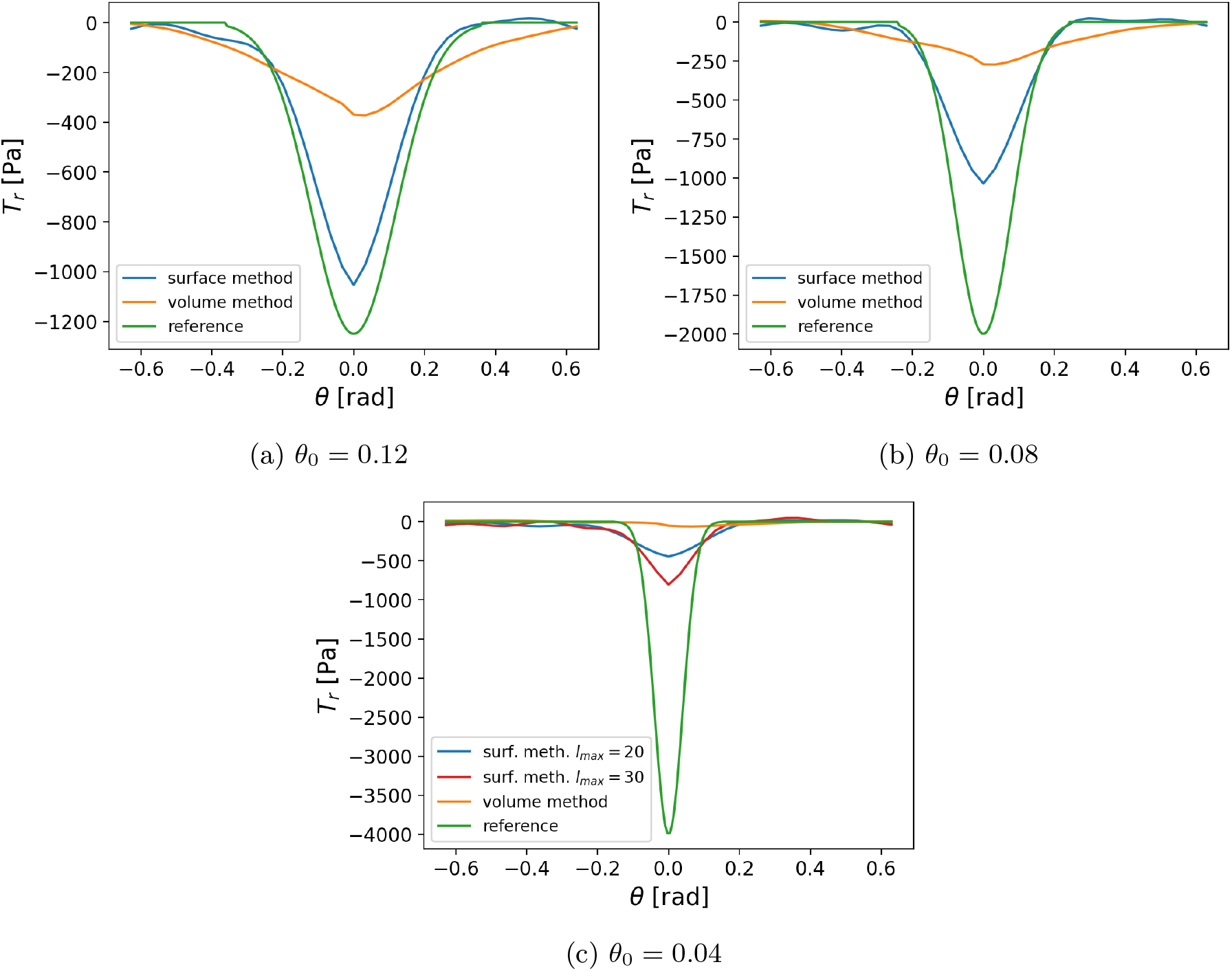
Performance decrease of both methods for highly localized indentation. The accuracy in recovering the reference profile strongly decreases with a decrease of the indentation width *θ*_0_. (a, b, c) Reference and recovered traction component *T*_*r*_ near the top pole of the sphere along y-axis for *x* = 0 (negative/positive *θ* corresponds to negative/positive y values). The width *θ*_0_ of the Gaussian indenter is varied in (a, b, c) as stated. The magnitude is varied accordingly to obtain similar indentation depths in all cases (≈ 20%). The results in Fig.2 correspond to the parameters used in (b).

**Figure S15.**
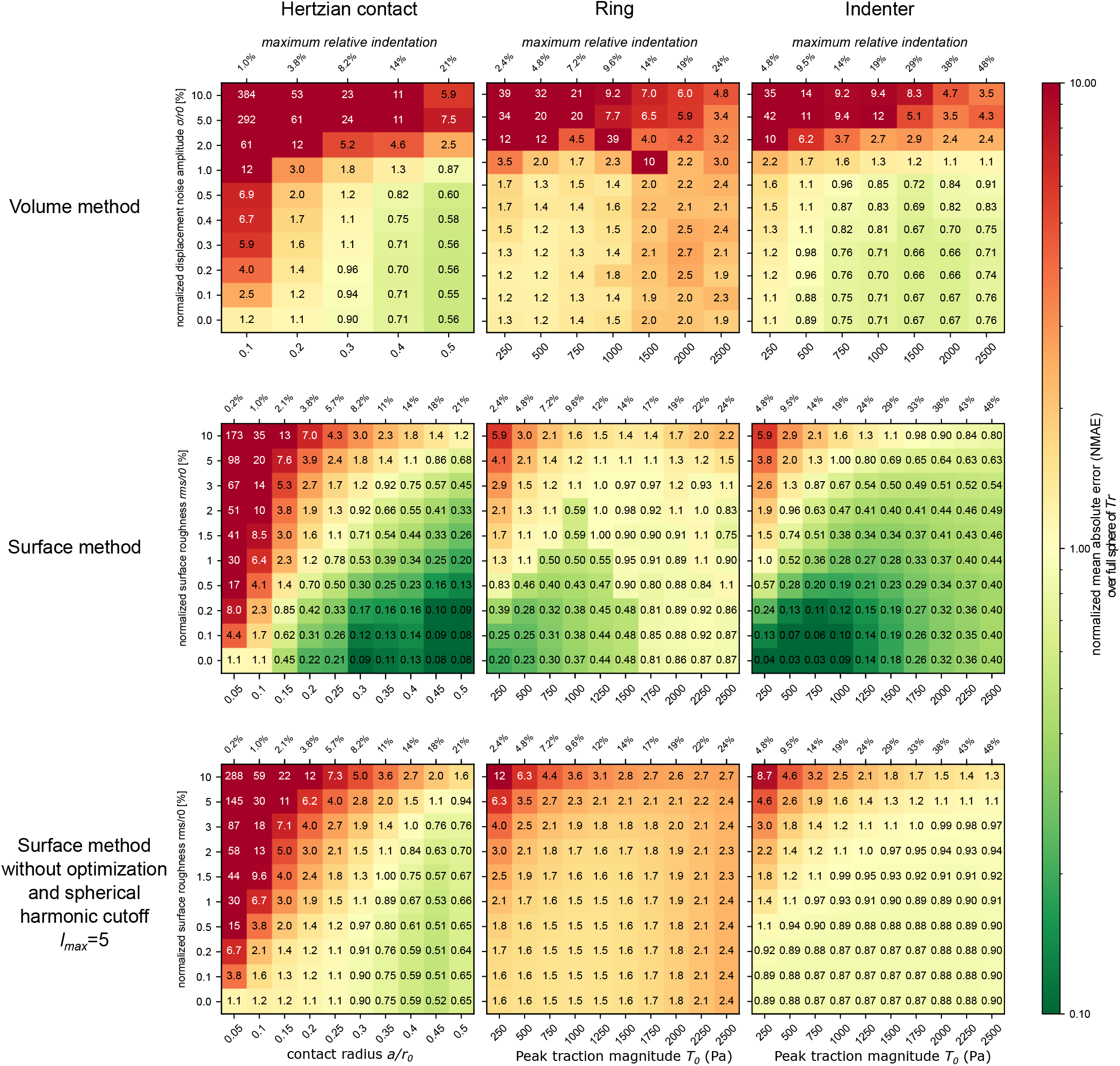
Full heatmap grids for method error comparison for full sphere evaluation of the NMAE of *T*_*r*_.

**Figure S16.**
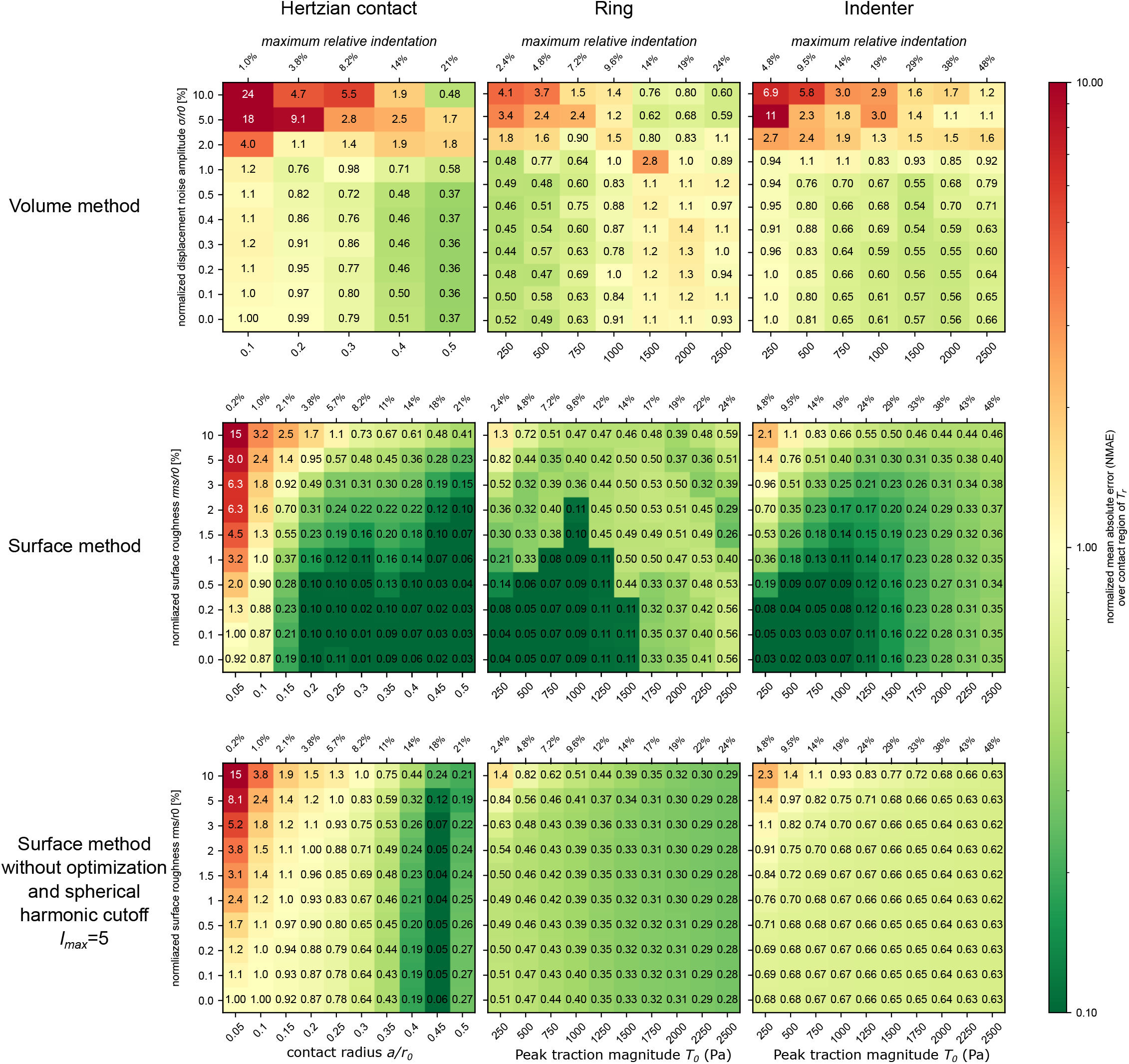
Heatmap grids for error evaluation restricted to dominant region. In Fig.3, the NMAE was evaluated across the whole sphere. Here we evaluate the NMAE in a region close to the region of maximum force application only. The same formula from Eq.(15) is used, however here we only sum over a smaller region Ω_0_. For the Hertzian contact and indenter profile, we choose a region near the poles close to the z-axis (i.e. *θ* ∈ Ω_0_ = [0°, 9°]∪ [171°, 180°]); for the ring profile close to the equator (Ω_0_ = [81°, 99°]).

**Figure S17.**
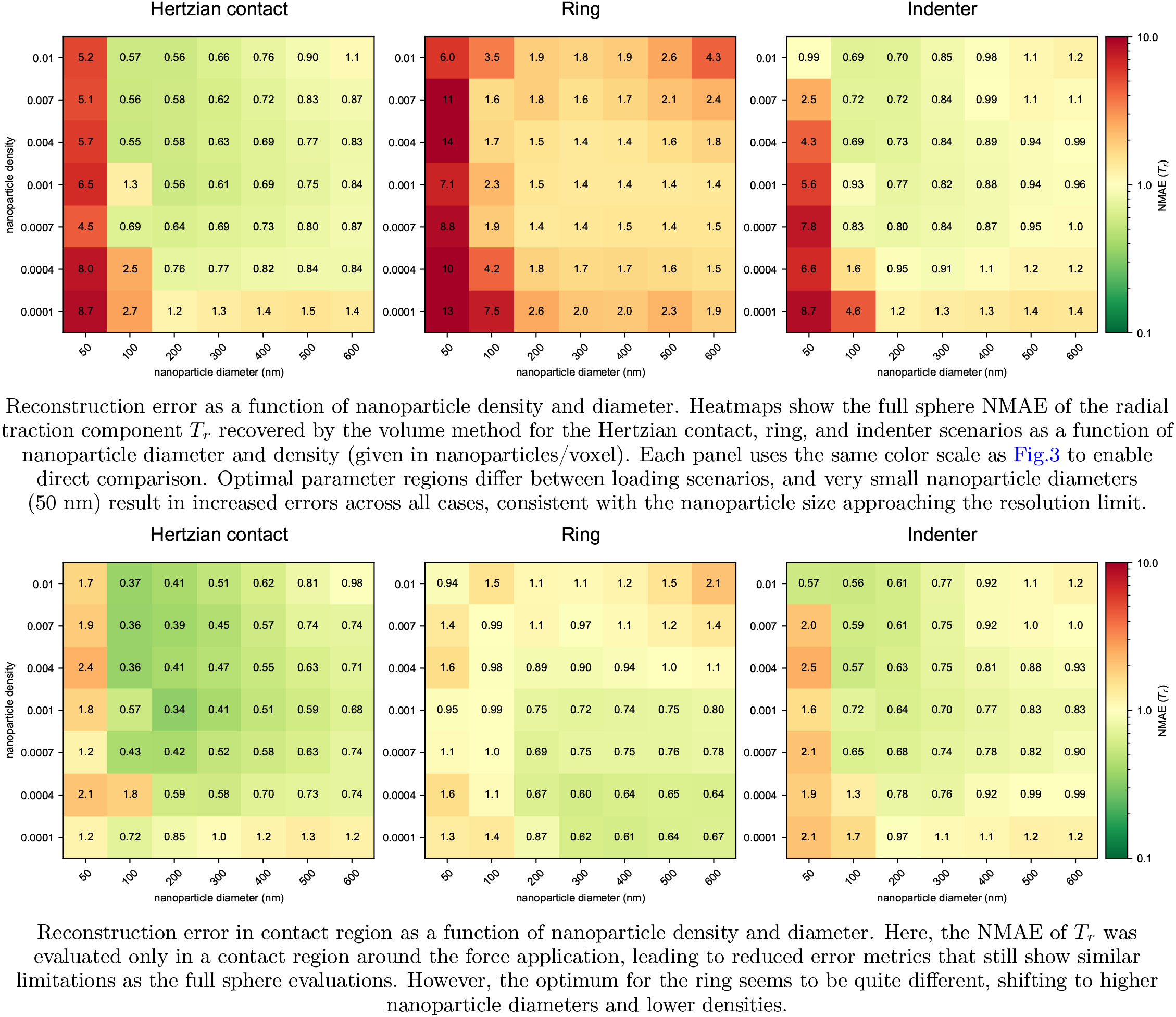
Effect of nanoparticle properties on volume method accuracy. Fig.3 shows that, under representative parameter settings, the volume method exhibits larger reconstruction errors than the surface method. Because the volume approach relies on displacement tracking of embedded fluorescent nanoparticles, its accuracy may depend on density and diameter of these fiducial markers. We therefore systematically varied these experimental parameters to assess whether improved marker-based displacement resolution reduces reconstruction errors, with full results for each configuration shown above. Importantly, we find that although suitable parameter choices reduce the error magnitude in specific regimes, even at the respective optimal conditions, the minimum reconstruction error of the volume method remains above that achieved by the surface method.

**Figure S18.**
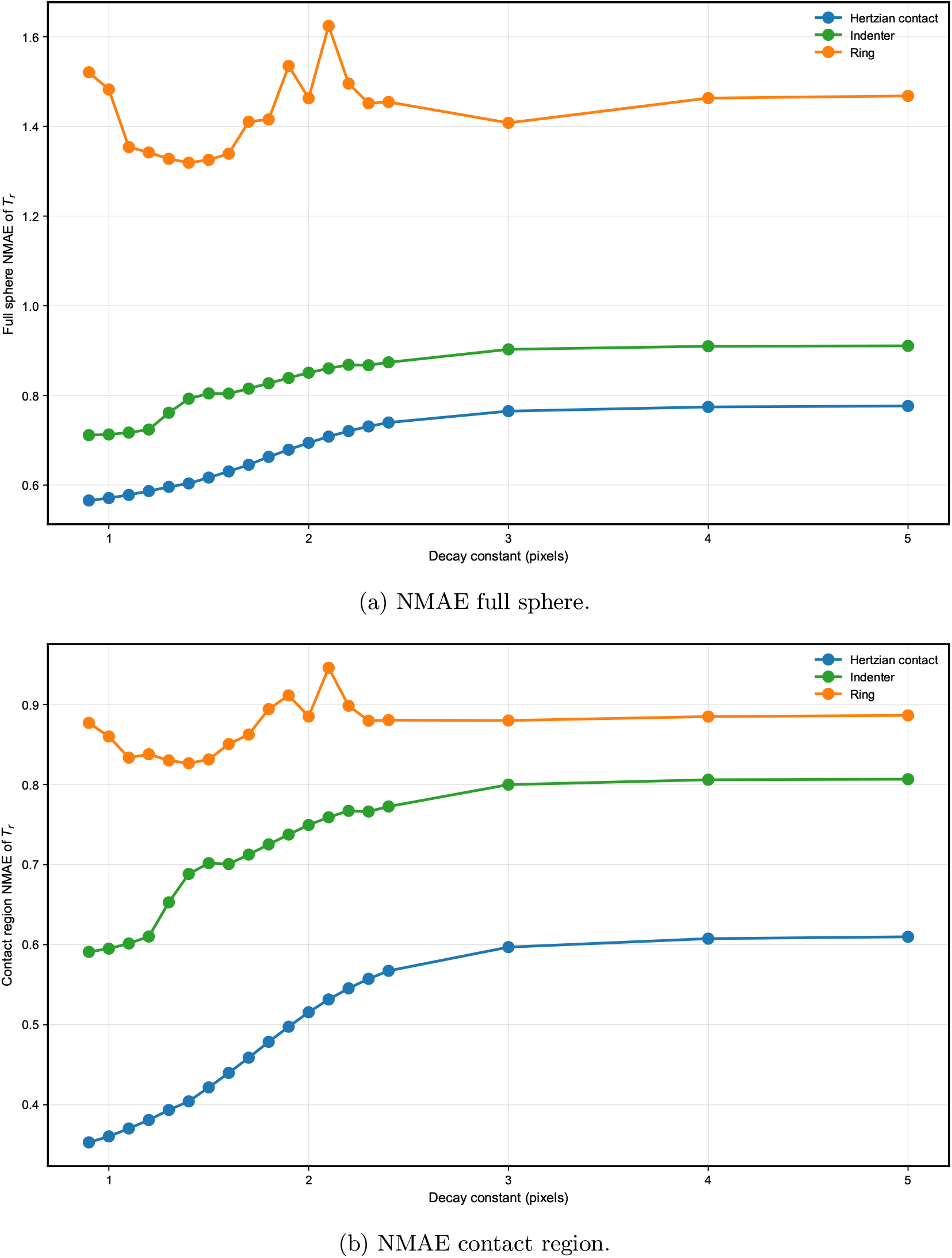
Simulation results for imaging artifact sensitivity via PSF decay constant sweep, mimicking different blurring levels. The decay constant *γ*_*d*_ controls the width of the PSF applied to the simulated nanoparticles in volume method image generation, with larger decay values representing heavier optical blur. The PSF as a function of the distance *r* from the center of the nanoparticle is defined as 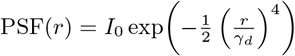, where *I*_0_ denotes the peak intensity in the center of the microparticle. Both *r* and *γ*_*d*_ are given in pixels. *γ*_*d*_ ranges from 0.9 (minimal blur, sharp PSF) to 5 (heavy blur, degraded optical conditions). Standard analyses in this work used a value of 1. NMAE of *T*_*r*_ is shown as a function of PSF decay constant, evaluated either over the full sphere (a) or only within the contact region (b), the latter isolating the regions where traction is applied and therefore revealing sensitivity to local force features. The analysis quantifies errors in reconstructing the prescribed traction profiles under increasing imaging blur and reveals scenario-dependent robustness. Localized contact geometries, including Hertzian contact and indenter cases, show substantial error degradation with increasing PSF blur, with an approximately 30% error increase from decay 0.9 to 5, reflecting the requirement for sharp image definition to accurately resolve localized contact boundaries for particle correlation. In contrast, the distributed stress gradient for the ring indenter remains robust to PSF blur, with only approximately 5% error variation across the decay range. These results suggest that contact geometry type fundamentally determines sensitivity to imaging degradation.

**Figure S19.**
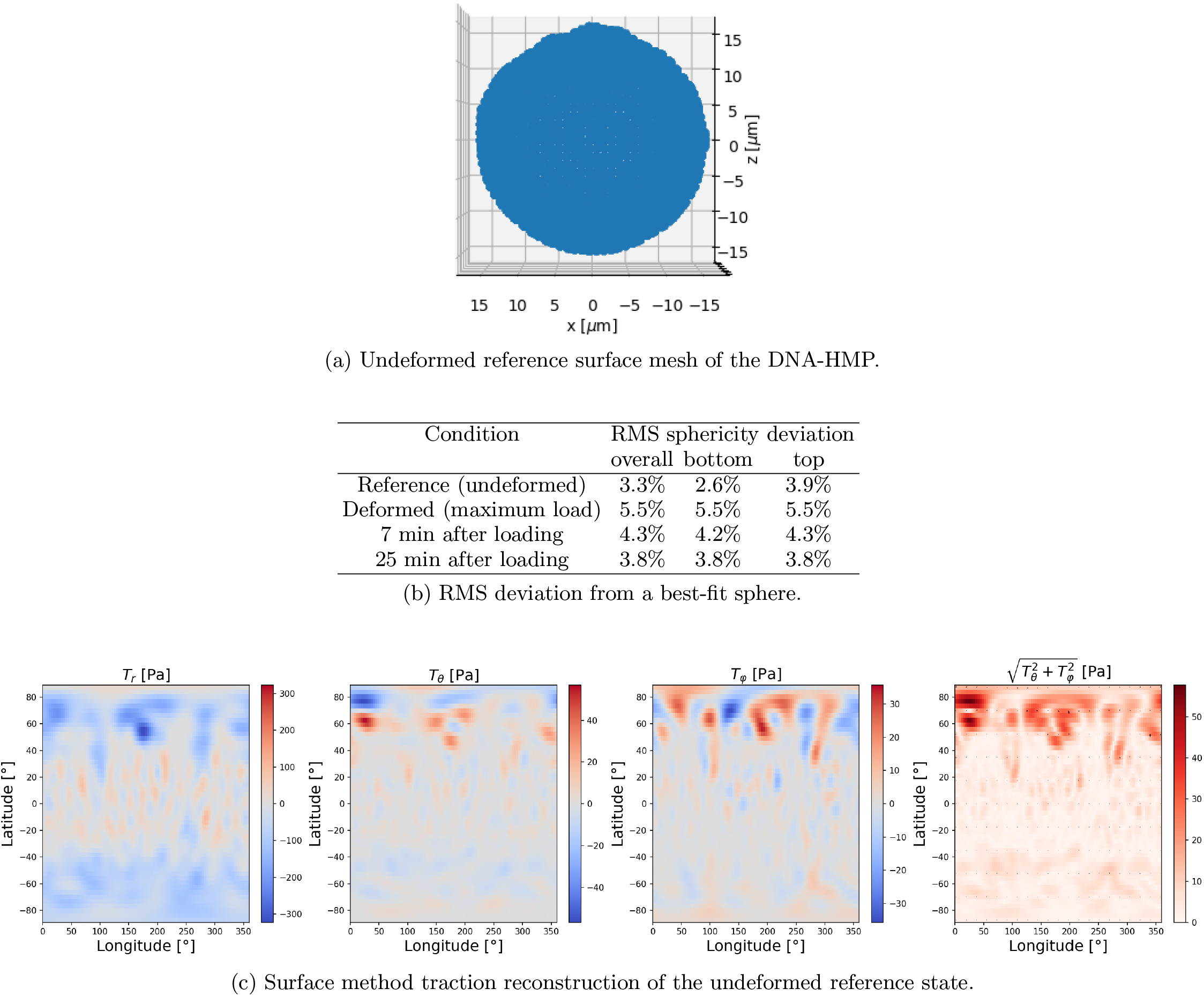
Quantification of sphericity deviation and residual tractions of DNA-HMP reference configurations. (a) Surface point cloud of the undeformed reference configuration of the DNA-HMP analyzed in the main text. The reference configuration remains overall spherical, supporting the undeformed spherical reference assumption. (b) RMS sphericity deviations from a best-fit sphere for different experimental conditions. After deformation, the RMS values return close to initial values, indicating recovery of spherical shape. Corresponds to Fig. S22(a–d). RMS sphericity deviation is defined as the root-mean-square of radial residuals normalized by fitted sphere radius. (c) Surface method traction reconstruction of the undeformed reference configuration, showing traction components *T*_*r*_, *T*_*θ*_, and *T*_*φ*_, and shear magnitude. Residual tractions are low compared to the deformed state, particularly in regions with better imaging quality. Minor residuals may arise from geometric asymmetry or imaging artifacts. These results support the validity of the undeformed spherical reference assumption for DNA-HMP traction reconstruction.

**Figure S20.**
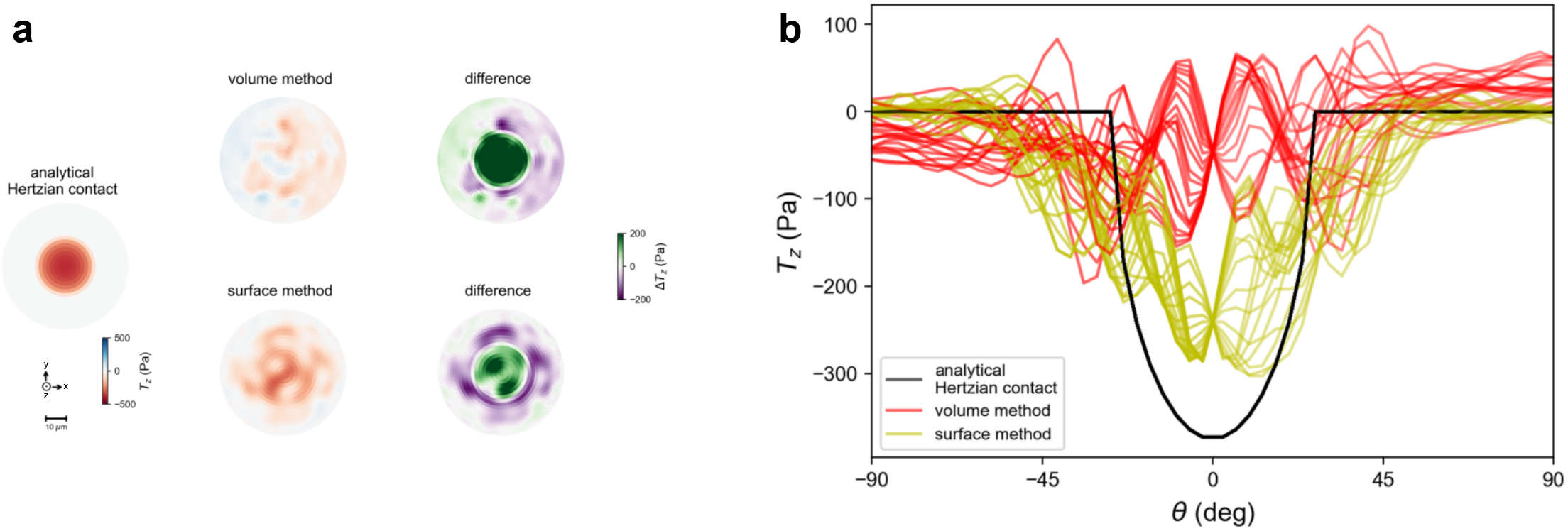
Analogous to Fig.5 (without the experimental setup in subfigures (a-d)), shown here are the reconstructed traction profiles of the top side of the same microparticle together with the same reference Hertzian contact (*a* = 0.45). (a) Analytical reference (left) and volume and surface method reconstructions (center) of *T*_*z*_, together with the deviation from the reference (right). (b) Superimposed cross-sections. The resulting tractions do not reflect the Hertzian contact as well as the reconstructed bottom side profile in Fig.5(e,f), with the surface method capturing better correspondence. However, this might not be a shortcoming of the methods themselves: Due to our experimental setup, the deformation pattern at the top side where the glass slide was pushing down, as well as the image quality, was less clear, because the glass slide could move (while the bottom is fixed). Signal quality and resolution therefore were decreased. This is visualized (particularly for the surface method) in S21, which might also point at increased deformation at the bottom. The cross-sections still show that the compression can be captured by the surface method, and since the NMAE was evaluated for both methods over both contact regions (top and bottom side), reconstruction quality at the bottom is better than reflected by these numbers, implying higher quality traction profile reconstructions are possible with improved experimental setups.

**Figure S21.**
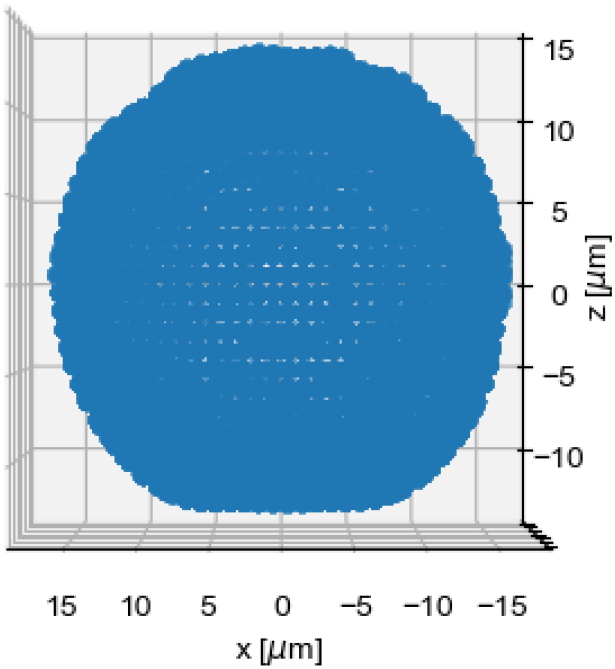
Set of points describing the microparticle surface of the DNA-HMP analyzed in Fig.5 and S20, obtained from using GeoV^47^ (after ImageJ^46^ preprocessing) on the microscope data of the Cy3-label in the DNA network. Noticeably, signal quality at the top seems decreased, and the deformation is less clearly visible (possibly also because the force application had a different contact radius on this side). Reasons for quality decline at the top might be that the glass slide is not fixed like the bottom, and optical effects leading to lower signal on top. While the set of points itself is only used in the surface method, these general experimental shortcomings could explain the worse top side reconstructions by both methods shown in Fig.S20, Fig.S23(b), and Fig.S24(b, d).

**Figure S22.**
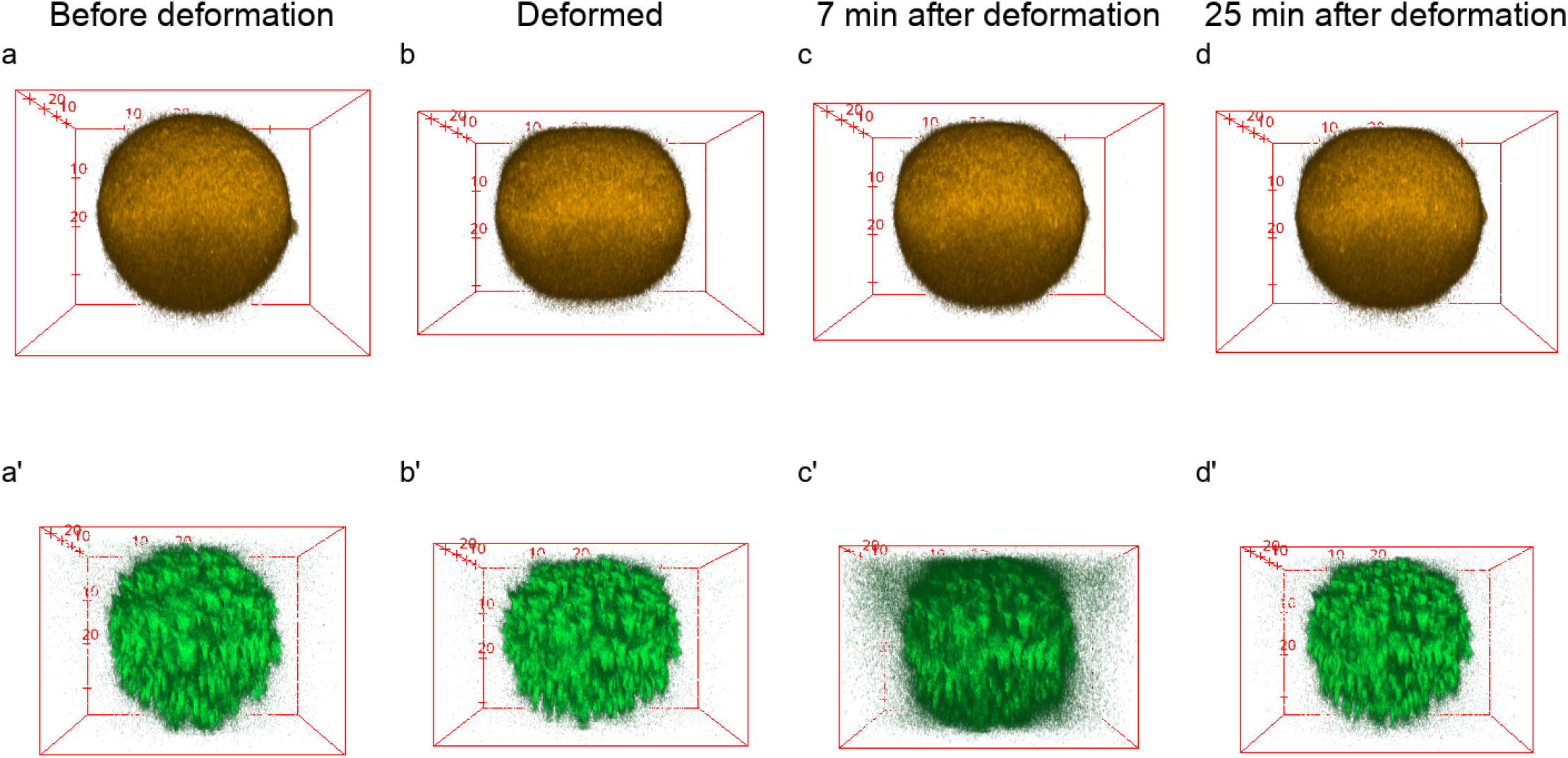
Microscope images of the DNA-HMP analyzed in the main text (Fig.5), visualized using 3D Viewer in ImageJ.^46^ Note the z-axis is pointing downwards here. First row (a, b, c, d) shows the Cy3 labels of the DNA network, used for the surface method reconstruction. Second row (a’, b’, c’, d’) shows the nanoparticles tracked in the volume method. From left to right, the columns represent the images taken before the deformation from the top by a weight, the deformed state, and two relaxation images captured after the weight was lifted (7 min and 25 min after). The volume method for the main text analysis used the relaxation of 25 min as reference image.

**Figure S23.**
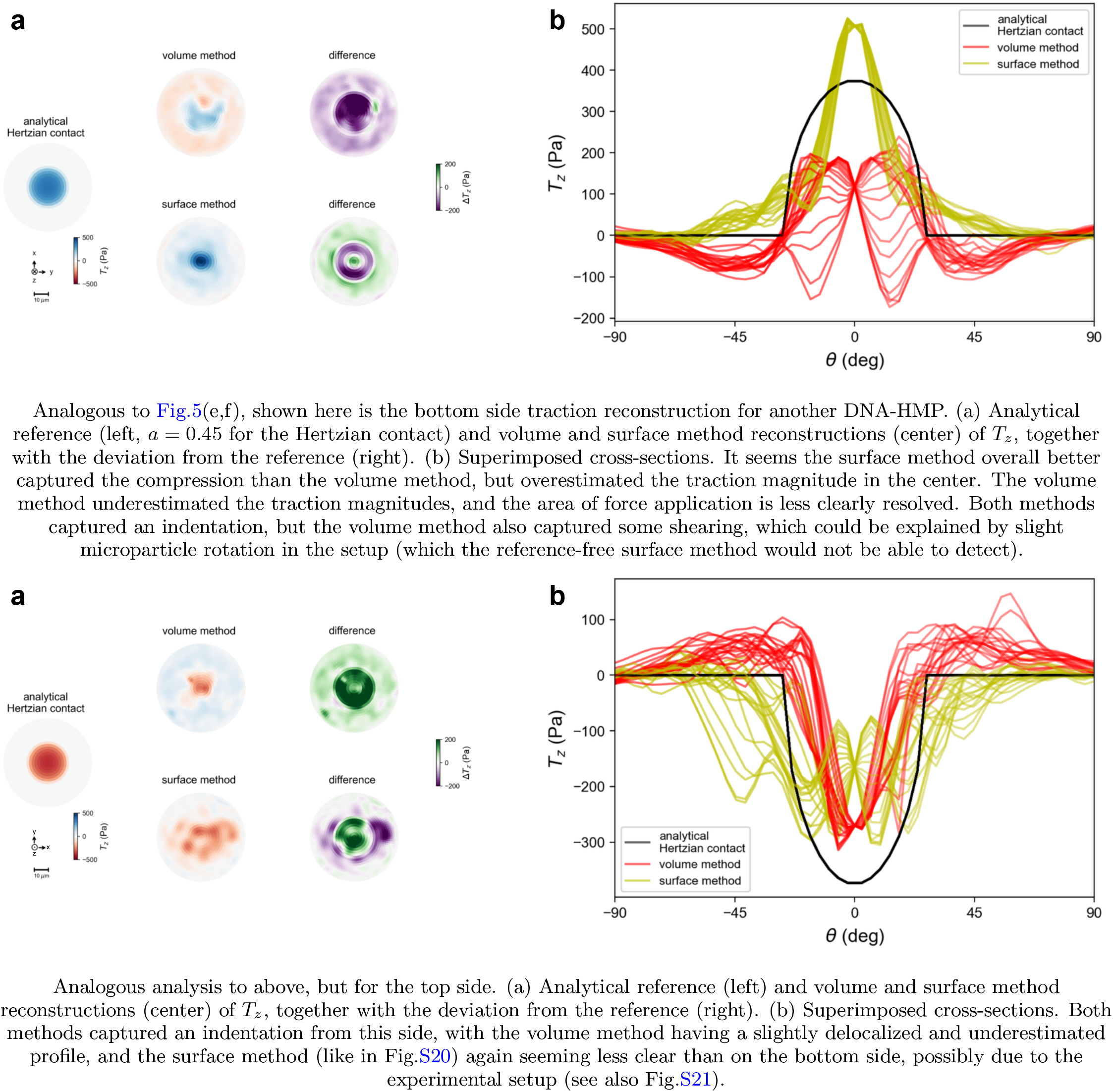
Traction reconstruction results for both surface and volume method for another DNA-HMP.

**Figure S24.**
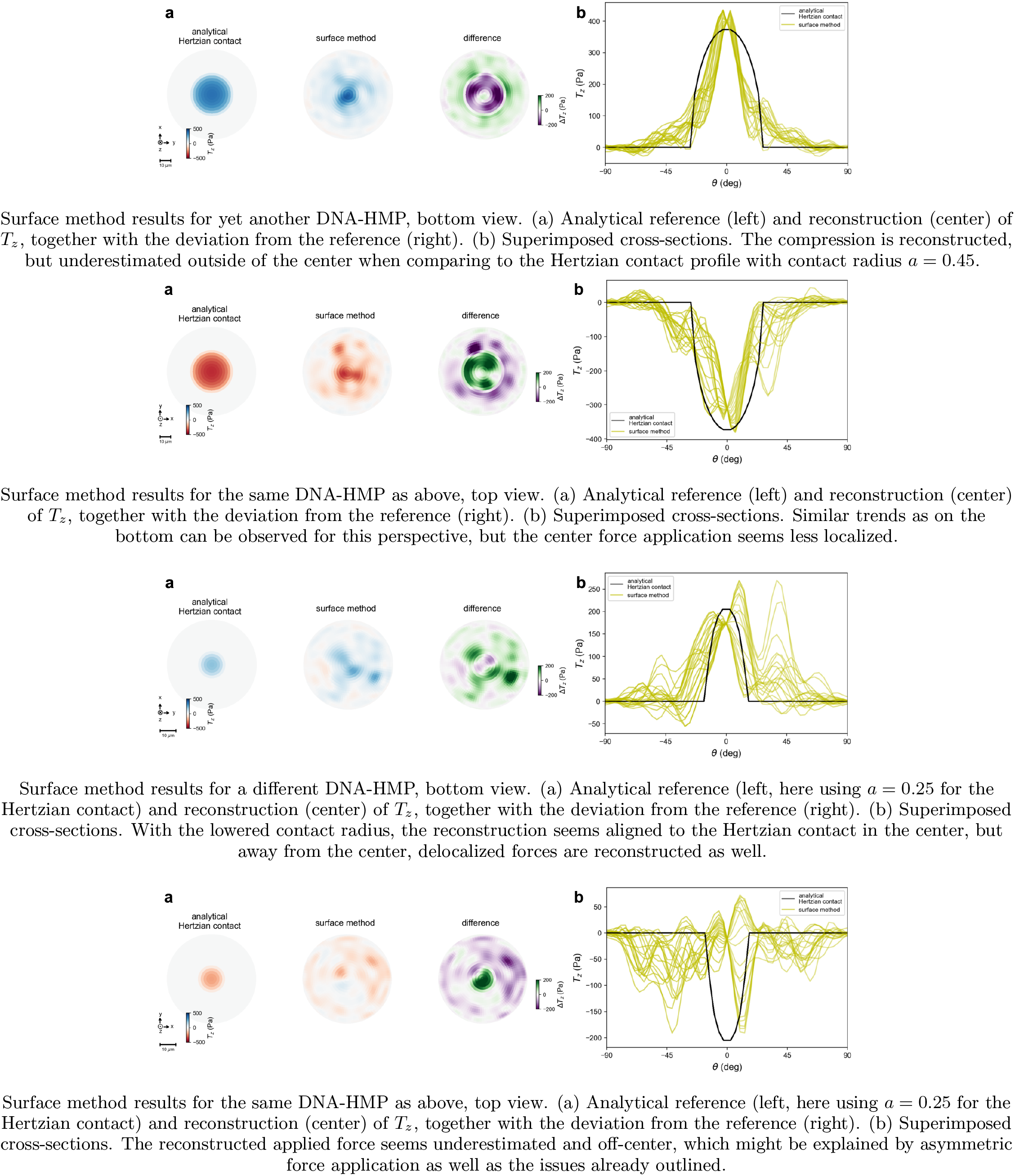
Surface method results for two more DNA-HMPs.

**Table S1.** Standard simulation and analysis parameters for volume and surface method, used in this work if not stated otherwise. The lower resolution in the z-direction for the volume method resembles the lower resolution typically obtained by z-stack scanning in experiments.

| volume method |  | surface method |  |
| --- | --- | --- | --- |
| parameter | value | parameter | value |
| microparticle properties: |  | microparticle properties: |  |
| radius $r_0$ | 12 $\mu\text{m}$ | radius $r_0$ | 12 $\mu\text{m}$ |
| elastic modulus $E$ | 1500 Pa | elastic modulus $E$ | 1500 Pa |
| Poisson's ratio $\nu$ | 0.4 | Poisson's ratio $\nu$ | 0.4 |
| resolution | 120 nm(x,y); 240 nm (z) | cutoff $n_{\text{max}}$ | 50 |
| nanoparticle density $\rho_n$ | 0.002 nanoparticles/voxel | GLQ mesh order $l_{\text{max}}^*$ | 50 |
| nanoparticle radius $r_n$ | 100 nm | deformed surface simulation parameters: | |
| cutoff $n_{\text{max}}$ | 50 | number of surface (Fibonacci) points $N$ | 2000 |
| displacement noise $\sigma_{\text{disp}}$ | 0.17 px | roughness amplitude $\sigma_r$ (relative to $r_0$) | 0.02 |
| padding factor $p$ | 1.28 | power-law exponent $\beta$ | 3 |
| resulting picture size | 256x256x128<sup>b</sup> | minimum SH degree $l_{\text{min}}$ | 2 |
| output spacing FIDVC | 4 | minimization parameters: |  |
| gradient method | optimal-5 | functional prefactors $\{\alpha, \beta, \gamma\}$ | $\{1, 1, 0.3\}^c$ |
| convergence criteria | default | # iterations/period | 10 |
| evaluation radius $r_e$ | 0.8 $r_0$ | # periods $n_p$ | 15 |
| | | SH cutoff $l_{\text{max}}$ | 20 |

**Table S2.**
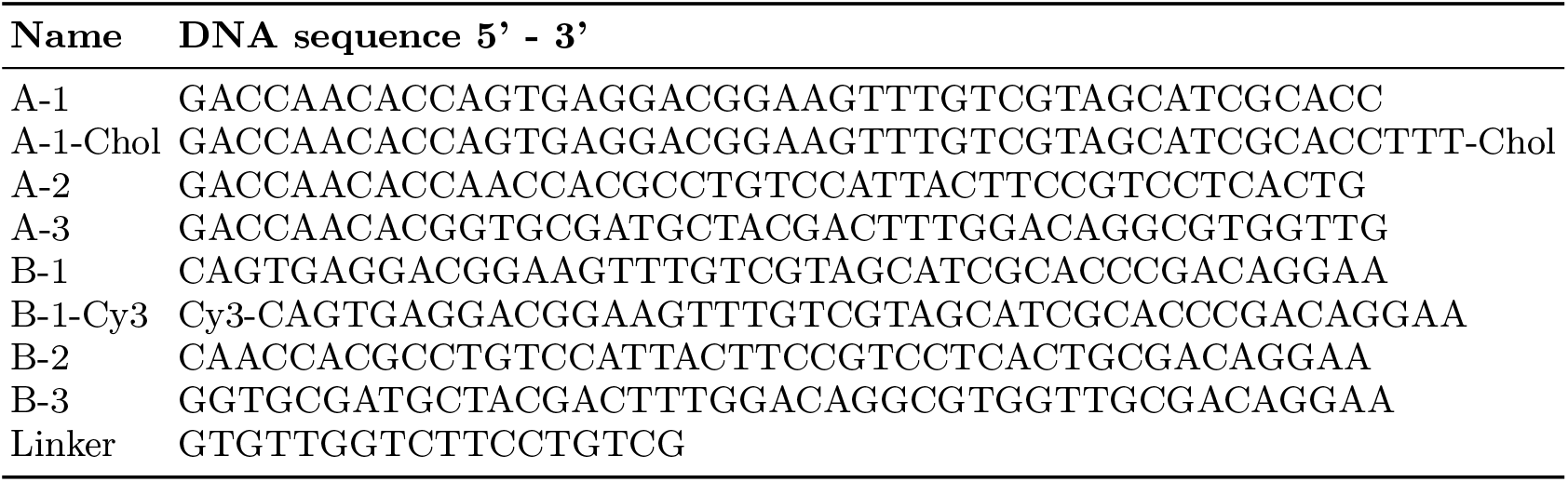
DNA sequences used for the formation of the DNA-HMPs. The fluorescent label cyanine 3 is abbreviated as Cy3. Cholesterol is abbreviated as Chol.

### Note S1: SH decomposition of displacement and traction

The decomposition described in this step is based on the work of Wang et al. in 2019.^36^. To evaluate the functional quickly, it is necessary to obtain a fast way to convert from 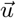 to 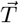, as the terms in the functional have an explicit dependence on 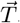. The idea here is to choose a basis for the displacement function space where each basis vector 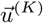 corresponds to a basis vector 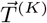, i.e. that if a displacement field is given as a superposition

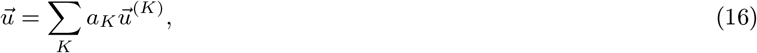

the corresponding traction is given by

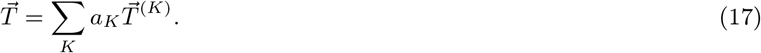

Such a basis can be constructed by expressing the quantities and thus also the basis vectors in terms of spherical harmonics (SH), as explained in the following: We start with the Papkovich-Neuber ansatz (see Eq.(9)). Using a potential ansatz, the general harmonic solutions are then given by

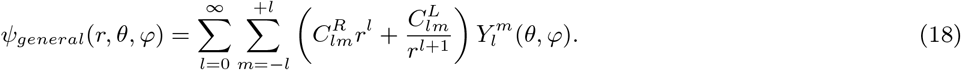

The part of the radial solutions with negative exponent in *r* lead to solutions for 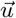 diverging at the origin, so we set 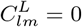. As we want to get a set of basis vectors, we use that the spherical harmonics *Y*^*m*^ provide an orthonormal basis for functions of the angles (*θ, φ*) projecting onto ℝ or ℂ and use them as a basis. As *ψ*_*k*_ (and later *u*_*k*_ and *T*_*k*_) with *k* ∈ [*x, y, z*] contain three components each, we need a full set of SH for each component *k* separately to cover the full solution space. Taking this into account, we choose a set of 3 numbers *K* = (*k, l, m*) to denote our basis vectors as

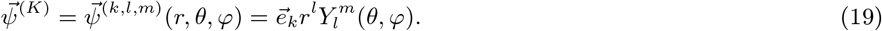

Switching to index notation, we now plug the potential basis vectors into Eq.(9) and obtain^1^

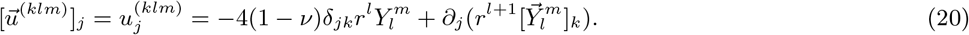

Here we have set *ψ*_0_ = 0, as we can recreate any *ψ*_0_ with a suitable choice of coefficients in the 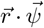-part in the same bracket.^36^ We also introduced the first vector spherical harmonics (VSH1) 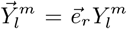. The single components *k* of the VSH1 can be decomposed into a superposition of scalar SH with coefficients *Q*:^36^

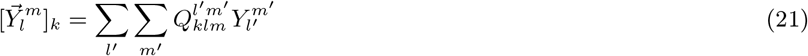

Plugging this into Eq.(20) and executing the derivative, we get

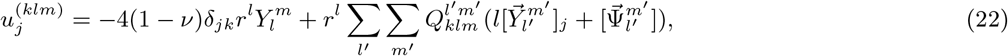

introducing the second vector spherical harmonics (VSH2) 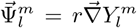. Similarly to the VSH1, their components can be decomposed into SH using a second set of coefficients *P*:

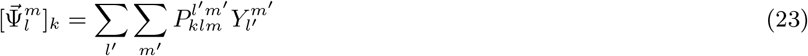

Replacing the VHS2 in Eq.(22) with their SH decomposition, and doing so as well for the VSH1 once more yields:

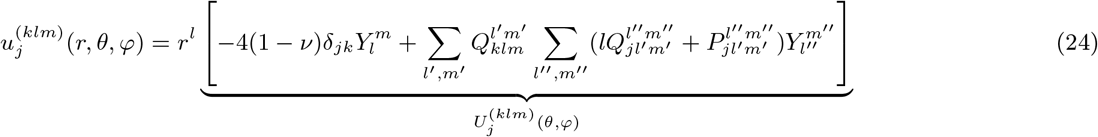

Here we factored out the polynomial in *r*. Identifying the *r*-independent part 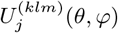, we can now further decompose it. Explicitly factoring out the sum 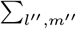 in the second term and considering the first term, we can separate the SH and define the resulting coefficients as 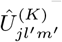:

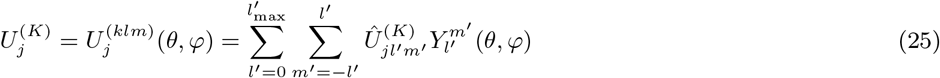

As a result, each basis vector is now fully described by a decomposition into spherical harmonics. Here we introduced a cutoff *l*_max_, which can be set to favor either computational speed (low *l*_max_) or accuracy (high *l*_max_). For the next step, we use the spherical versions of the strain and stress tensor:^49^

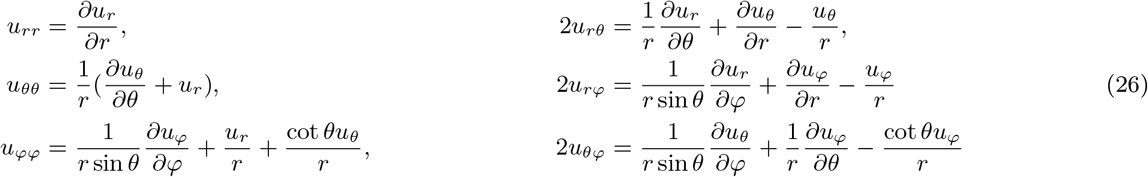

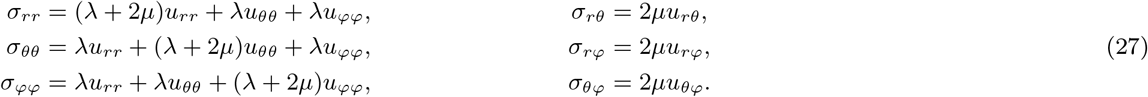

Evaluating Eq.(4), which can also be rewritten in terms of spherical harmonics, we obtain a similar separation and decomposition for the traction

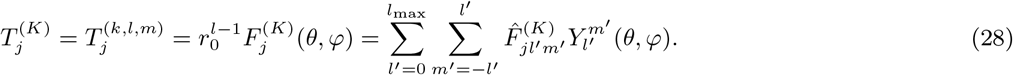

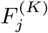 and 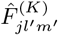 denote the *r*-independent part of the traction and its decomposition coefficients for the respective basis vectors.

The coefficients 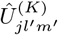 and 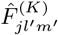 are fixed for a given set of material parameters (*E, ν, r*_0_), as the relations between the SH and VSH *Q* and *P* are also constant. Thus, they can easily be precomputed and used again in each iteration of the minimization.

### Note S2: Vector representation of the elastic energy

The subsequent procedure was introduced in the github package SHElastic.^50^ The general package has been developed for the publication describing the SH decomposition of the previous step.^36^ Example 06 of the package is the specific implementation for the surface method,^29^ including suitable vector representations and reformulations of *f*, which we will present in the following.

The definitions Eq.(25) and Eq.(28) allow us to easily represent the full displacement field as a vector 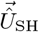, whose entries are the coefficients of the corresponding spherical harmonic (running from 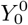 to 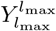 for the three displacement field components *u*_*x*_, *u*_*y*_, *u*_*z*_ successively).The vector representation also allows us to rewrite the functional in such a way that we can efficiently perform the minimization.

We also need a vector representation for the traction in position space to evaluate the *E*_*res*_ term easily. To do so, we choose a set of N points on the surface, characterized by their corresponding angles (*φ*_*i*_, *θ*_*i*_). We choose a vector 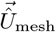, whose entries simply describe the displacement values at the respective points *i* (The first *i* components describe *u*_*x*_ at the respective point, components *N* + *i* describe *u*_*y*_, etc.).

As the set of points, we use a GLQ mesh (Gauss-Legendre quadrature mesh, characterized by order *l*^*\**^, which we choose to be the same as the SH, i.e. *l*^*\**^ = *l*_max_). This mesh is used as it facilitates a fast conversion between the mesh and the SH coefficient representation of the displacement field: It can be implemented as a matrix multiplication with a precomputed complex conversion matrix **S** (performed using *SHTools*^51^):

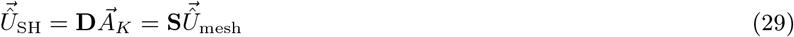

This intermediate definition resembles Eq.(16) in vector representation, where the coefficients *a*_*K*_ are represented as one vector 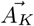. The entries of the matrix **D** consist of the coefficients 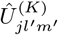 to relate the contributions of the corresponding basis vector *a*_*K*_ to the SH modes. For the traction, we similarly find:

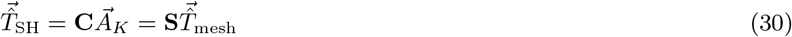

We can relate the traction and displacement via 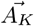 in the two equations, leading to:

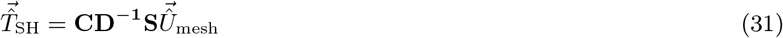

Plugging Eq.(30) into Eq.(31), we can directly relate 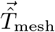 and 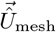:

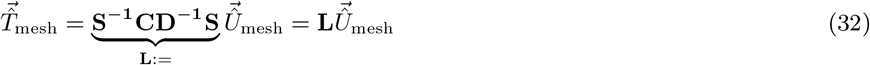

Note that the vectors with the hat 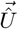 describe the displacement field 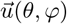 but other than 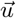, they are not vector fields themselves, but a vector representation of the field. The same holds true for the respective traction vectors.

With these definitions, we can rewrite the three different parts of our functional *f*. Using the definitions above, we also reformulate them to depend explicitly depend on 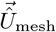, as we need to perform derivatives with respect to it later.

1. For *R*, we introduce a matrix **P** that selects the points of the mesh on the traction-free region and also contains weighing factors to compensate for the higher density of points near the poles for the GLQ mesh, which consists of *N* = (*l* + 1)(2*l* + 1) points, and obtain

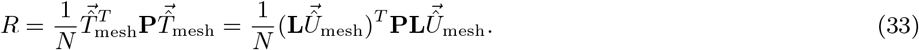
2. For *E*_el_, instead of integrating across the surface, in the SH representation we can evaluate the integral with the orthogonality relation of the SH: 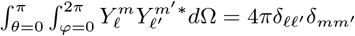. In the vector representation, this leads to

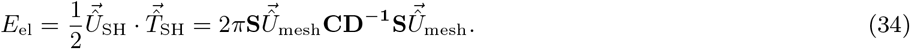
3. We can now also define the high-frequency penalty in a meaningful way. For this, we use the matrix **Q** that introduces a damping coefficient for higher order traction modes,^29^ which correspond to high spatial frequency components.

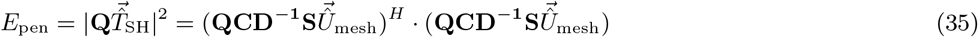

(…)^*H*^ denotes the Hermitian transpose to ensure we get the absolute value, as **S** contains complex elements.

### Note S3: Minimization

In order to start the iteration, we need an initial guess for the displacement field. The easiest choice incorporating the measured shape of the deformed sphere is to assume the displacement is only happening in radial direction. As we know the surface shape of the sphere, we can determine the radial change for each point on the surface: Using linear interpolation, we define a function 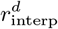 that, after calculating the volumetric central point 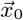 of the deformed sphere, assigns a radius value for the whole measured surface, parameterized by *θ* and *φ*. The radial displacement is then simply given as

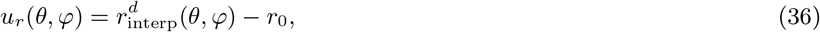

where *r*_0_ denotes the reference radius of the undeformed sphere. However, obtaining the true reference radius is unfortunately impossible without a reference picture. In theory, the sphere could have had any possible radius before, it could even have been compressed strongly and there would be no way of recovering this information from the single picture. The easiest solution for this is to compute the volume of the sphere and use the radius of a sphere of the same volume, given via 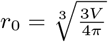.

With all the earlier stated advantages that taking only one picture brings, this is a major disadvantage of this method compared to the volume method: We basically assume that the material is incompressible and lose all information about isotropic normal traction components (pressure) acting on the sphere. We also assume the reference shape to be a perfect sphere, introducing another error source as most produced microparticle will probably slightly deviate from this shape. For the volume method, this is accounted for with the reference picture as well, containing any deviations from the spherical shape.

As stated before, we will use a GLQ mesh as the set of points *i* to perform the minimization on, each point being characterized by its position (*θ*_*i*_, *φ*_*i*_). As the surface shape is fixed, this limits the ways in which we can update the displacement during the minimization. To ensure that this boundary condition is always obtained, we choose the angular surface shifts 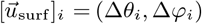 of the points *i* as the degrees of freedom that are updated throughout the minimization process. The surface shifts denote the change in solid angle from the reference mesh point. In this way, the corresponding radius can be determined from the interpolation function via 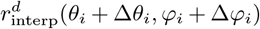 and the boundary condition is always fulfilled. For the initial guess of the displacement field being only in the radial direction, this corresponds to (Δ*θ*_*i*_, Δ*φ*_*i*_) = (0, 0) for every point *i*.

To update the field, we need the full Jacobian of the functional *f* with regard to the vector including all surface shifts 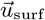:

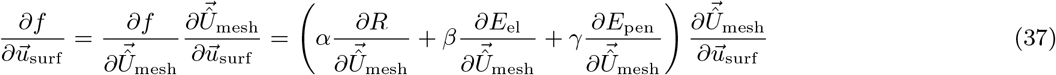

The entries of the Jacobian indicate how the functional changes with a change of the single surface shifts, representing a high-dimensional derivative. The derivatives of the three terms (*R, E*_el_, *E*_pen_) are given explicitly in example 6 of the corresponding GitHub package of the surface method.^50^ To evaluate the second term 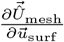, we need to account for the change of the interpolation function 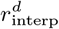 with regard to the surface shifts (Δ*θ*_*i*_, Δ*φ*_*i*_), as this directly determines the change in *u*_*r*_. We approximate the derivative by calculating the 3-point gradient in a small environment around the respective position (*ϵ* = 10^*−*5^) for each point *i*:

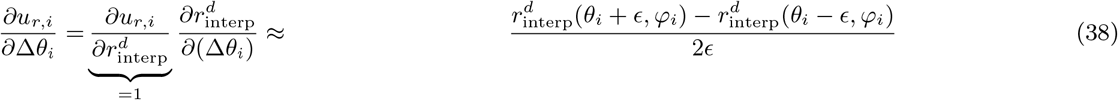

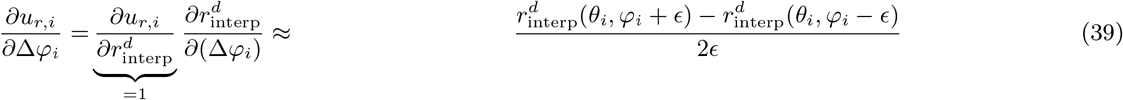

As they are given in radians and we can assume them to be small, the tangential displacement components are linear in the small angular shifts after scaling with radius; therefore the corresponding Jacobian entries are:

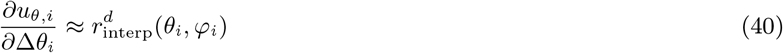

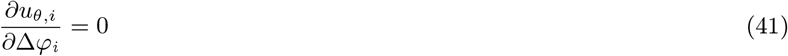

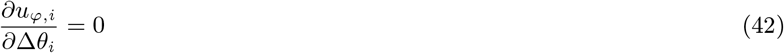

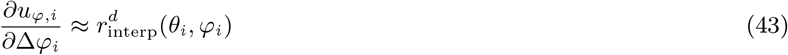

Transferring these to Cartesian coordinates yields the needed relation.

The minimization itself is performed using the conjugate-gradient method.^52^ As specific implementation, we use the *minimize* function of the *scipy*.*optimize* package.^53^ The surface shifts are updated until we reach the maximum number of periods *n*_*p*_.

### Note S4: Traction profile definitions

#### Sphere-wall contact (Hertzian contact profile)

The Hertzian contact profile describes the contact between an elastic sphere and an elastic half-space (characterized by (*E*_0_, *ν*) and (*E*_hs_, *ν*_hs_) respectively). It can be used to model the contact between an elastic sphere and a wall and is often utilized in AFM setups to estimate the elastic modulus *E* for materials. For the limit of small strains, this problem was solved in 1882 by Heinrich Hertz.^54^ It is equivalent to the contact between a rigid wall and an elastic sphere with an effective elasticity modulus *E*^*\**^:

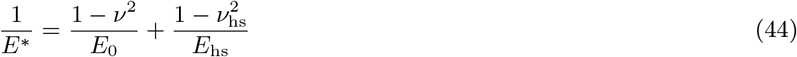

The resulting traction field on the sphere is originally parallel to the z-direction,^55^ decomposing it via *T*_*r*_ = *T*_*z*_ cos(*θ*) and |*T*_*θ*_| = |*T*_*z*_ sin(*θ*)| yields:

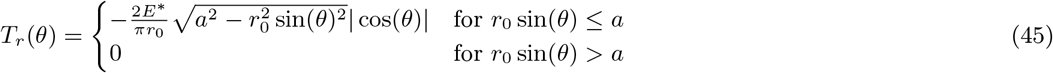

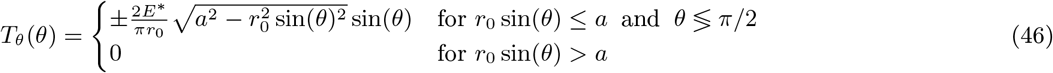

*a* describes the radius of the area where the sphere touches the wall, demonstrated in Fig.2. It is connected to the total applied force *F* via

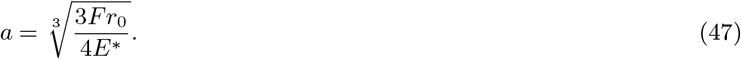

#### Gaussian indenter

One example of a more localized profile is that of an indenter pushing on the top and bottom of the sphere. We model this scenario by applying a localized Gaussian distributed traction profile of width *θ*_0_, resembling the standard deviation. We parametrize the Gaussian directly via *θ*:

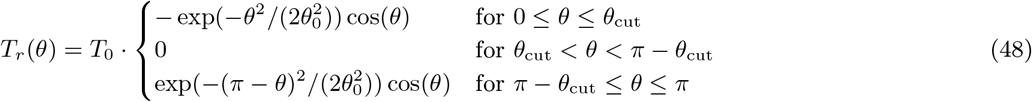

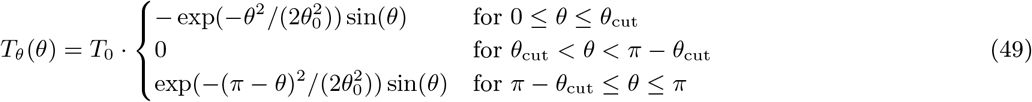

We introduced a cutoff angle *θ*_cut_, which is generally 3 times the size of *θ*_0_. In this way, we do not unnecessarily shrink the traction free-region for the surface method, but still get a smooth profile. *θ*_0_ and *θ*_cut_ are usually small. Considering the small angle approximation, we can assume that *θ*_0_ corresponds to a contact/cutoff radius in terms of *r*_0_. For example, if *θ*_0_ = 0.08, the width of the Gaussian corresponds to 8% of the radius of the sphere (≈ 1*µ*m for *r*_0_ = 12*µ*m in the simulations).

#### Gaussian ring

Another localized scenario is that of a ring-shaped traction profile of a certain width around the equator, as visualized in Fig.2(b). The motivation to examine this scenario is that in the original work of the surface method, a similar traction profile was exerted on a microsphere when being subjected to phagocytosis.^29^ We choose a Gaussian traction distribution with similar parameters to the indenter profile, centered around the equator at *π/*2:

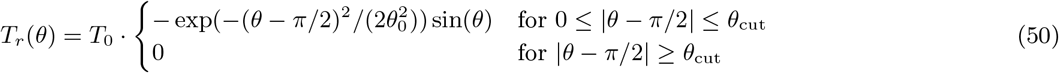

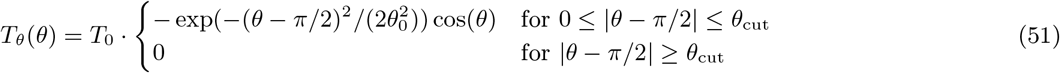

## Footnotes

1 We want to point out the difference between the indices *j* and *k* at this point to avoid confusion: *k* is part of the indication of the basis vectors (next to *l* and *m*). *j* ∈ [*x, y, z*] specifies the components of that basis vector. For the initial definition of the basis vectors in Eq.(19) with 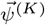, there is no use in distinguishing the two: As they are defined using 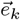, only the component *j* = *k* is non-vanishing. However, this is not necessarily true for the other basis vectors like 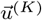, containing derivatives of 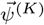 as the ∂_*j*_ in Eq.(20). Hence, in this and subsequent equations, the additional index *j* is necessary.

2 This picture size is very suitable for the FIDVC, as the sizes in all directions are powers of 2: Due to the window halving steps, other resolutions can cause asymmetric outputs of the FIDVC, which need to be accounted for in post-processing, unnecessarily complicating the evaluation procedure. The padding factor *p*, describing how much space we add around the sphere was chosen to obtain this resolution (*p* = 1 corresponds to the width of the picture being equal to the diameter of the sphere 2*r*_0_). This should also be taken into consideration by experimentalists when using this algorithm. The picture dimensions do not necessarily need to be an exact power of 2, it could also be a multiple of powers of 2, depending on the desired resolution. Choosing it to be a multiple of 32 is usually sufficient.

3 These parameters were used in the original publication and shown to provide reasonable results in extensive numerical analyses.^29^ Technically, they also resolve unit discrepancies between the energy functional terms.

## Notes

https://github.com/brewburgr/MPTFM

https://doi.org/10.11588/DATA/KWOD5A

### Competing Interest Statement

The authors have declared no competing interest.

### Summary of Updates

Fig. 6 with a nonlinear analysis and supplemental Figs. S6, S18 and S19 are new.

